# Spatial co-expression and cell-cell communication inference from spatially resolved transcriptomics with CONCISE

**DOI:** 10.64898/2026.06.22.733860

**Authors:** Jia Zhao, Xinning Shan, Gefei Wang, Tinyi Chu, Chen Lin, Rui Chang, Hongyu Zhao

## Abstract

Cell-cell communication is fundamental to tissue organization, homeostasis, and disease progression. Recent advances in spatial transcriptomics provide unprecedented opportunities to systematically characterize ligand-receptor interactions directly within intact tissues. However, robust inference of spatial ligand-receptor interactions remains challenging because intrinsic features of spatial transcriptomics data, including spatial autocorrelation, variation in total molecular counts, and measurement errors, can induce spurious spatial co-expression and lead to inflated false-positive results. Most existing methods do not adequately account for these confounding factors, limiting the reliability of inferred cellular communication. Here, we present CONCISE, a statistical method for spatially constrained co-expression and ligand-receptor interaction inference that jointly models spatial autocorrelation, variation in total molecular counts, measurement errors, and spatial proximity constraints. CONCISE combines efficient moment-based parameter estimation with analytical hypothesis testing, enabling fast and statistically rigorous inference without restrictive distributional assumptions. Through extensive simulations, real-data permutation experiments, and biologically motivated negative-control analyses across different spatial transcriptomics platforms, we show that most existing methods presented inflated false-positive rates, whereas CONCISE achieved well-calibrated inference, robust false-positive control, and improved detection power. Application of CONCISE to high-resolution MERFISH and CosMx datasets from intestinal inflammation and non-small cell lung cancer further highlights its biological utility in disease contexts. CONCISE uncovered inflammation-associated fibroblast-specific interactions during intestinal inflammation and delineated complex tumor-immune and tumor-stromal signaling networks within the tumor microenvironment.

## Introduction

Cell-cell interactions are fundamental to the organization of multicellular systems and play essential roles in diverse biological processes [1, 2], including tissue development [3], homeostatic regulation [4], and responses to stress and disease [5, 6]. A major form of these interactions is cell-cell communication (CCC), mainly mediated by ligand-receptor interactions (LRIs). In this process, ligands released by one cell bind to receptors on another cell to activate signaling pathways that coordinate gene expression and cellular functions [1, 2]. Identifying context-dependent LRIs holds great potential to elucidate disease mechanisms and inform therapeutic target discovery.

Direct measurement of proteins mediating CCC remains technically challenging [1]. As a more accessible alternative, transcriptomic measurements of ligands and receptors from bulk and single-cell RNA sequencing (scRNA-seq) have been widely used to study CCC across biological contexts [7, 8, 9, 10, 11]. Many computational tools have been developed, including CellPhoneDB [12], SingleCellSignalR [13], CellChat [14], ICELLNET [15] and CytoTalk [16]. These methods infer LRIs by employing distinct scoring strategies to evaluate the co-expression of candidate ligands and receptors in transcriptomic datasets, leveraging curated LRI databases. Although these approaches have generated important biological insights, their non-spatial nature can lead to inflated false-positive rates, as CCC typically occurs between cells in close physical proximity [17].

Recent advances in spatial transcriptomics (ST) enable high-throughput, spatially resolved transcriptomic measurements in intact tissues, providing new opportunities for more accurate LRI inference. Many computational methods have been developed, aiming to identify LRIs from context-specific ST data. Most methods, including MERINGUE [18], SpatialDM [19], Copulacci [20] and LIANA+ [21], infer LRIs by evaluating spatial co-expression between candidate ligand-receptor (L-R) pairs using geospatial statistics. In contrast to non-spatial approaches, these methods incorporate spatial coordinates when assessing co-expression, reducing false-positive interactions that violate spatial proximity constraints. Methods such as SpaOTsc [22] and COMMOT [23] further refine interaction characterization among L-R pairs by estimating directional spatial signal flow from ligands to receptors with optimal transport. Despite these advances, accurate identification of LRIs from ST datasets remains limited by intrinsic characteristics of ST data that, if not properly modeled, can generate spurious spatial co-expression signals and consequently lead to false-positive discoveries.

First, spatial autocorrelation is an intrinsic feature of ST data, whereby nearby spatial locations tend to display similar expression levels [24, 25]. This property can substantially confound spatial co-expression analyses. As widely recognized in spatial statistics, assessing bivariate correlation in the presence of spatial autocorrelation can markedly underestimate uncertainty, often to varying degrees, thereby leading to spurious associations and inflated false-positive rates if spatial autocorrelation is not properly accounted for [26, 27, 28, 29]. Nevertheless, existing ST-based LRI inference methods incorporate spatial information primarily to restrict candidate LRIs to spatial nearby locations, but do not account for spatial autocorrelation in ligand and receptor expression themselves. In particular, SpatialDM assumes that gene expression is independent of spatial location in order to derive an analytical null distribution for LRI inference, whereas other methods rely on permutation-based tests for L-R co-expression, which implicitly depend on the same assumption. Consequently, existing LRI methods can be seriously confounded by varying levels of spatial autocorrelation in ligand and receptor expression.

Second, additional properties of ST count data further challenge LRI inference. ST data exhibit varying total molecular counts across spatial locations, and expression counts are sparse and noisy [30, 31, 32]. If the count-based nature of the data, variation in total molecular counts, and measurement errors are not properly modeled, LRI inference can be distorted. Similar challenges have been recognized in scRNA-seq analysis, where ignoring these factors can substantially confound downstream analyses including gene-gene co-expression inference [33, 34, 35]. However, most existing ST-based LRI methods still rely on CPM normalization to adjust for total molecular counts prior to spatially constrained co-expression inference. Despite its simplicity, this strategy can introduce artificial co-expression signals, leading to false-positive results. Moreover, existing approaches often do not model measurement errors in the data-generating process, and rely on restrictive distributional assumptions, increasing the risk of spurious results [34, 35]. Taken together, these limitations highlight the need for methods capable of jointly accounting for spatial autocorrelation, count nature of the data, varying total molecular counts and measurement errors when inferring LRIs from context-dependent ST data.

Here, we introduce CONCISE, a principled statistical method for co-expression and cell-cell communication inference from spatially-resolved transcriptomics. Through innovations in model and algorithm design, CONCISE addresses the aforementioned challenges within a unified statistical framework. Specifically, CONCISE introduces the following key methodological advances for spatial CCC analysis. First, CONCISE models the unobserved true spatial expression levels as latent variables and links them to the observed count data through a statistical measurement model. This explicitly accounts for the count-based nature of ST data, variation in total molecular counts, and measurement errors. Second, by treating the latent expression levels as spatial processes, CONCISE jointly models ligand and receptor expression through a spatial process model. This enables CONCISE to infer L-R co-expression under spatial proximity constraints while adaptively accounting for different levels of spatial autocorrelation in ligand and receptor expression, mitigating false-positive discoveries arising from spatial autocorrelation. Third, CONCISE proposes an efficient moment-based approach for inferring spatial LRIs. This approach avoids imposing a restrictive distributional family for the underlying expression levels and can flexibly accommodate the data-generating process. Importantly, with this approach, CONCISE explicitly derives the analytical null distribution, enabling fast and principled statistical testing of spatial LRIs under the proposed model, thereby facilitating efficient and reliable CCC analysis from ST datasets.

To evaluate the performance of CONCISE, we conducted comprehensive simulation and permutation studies based on real data. These analyses show that CONCISE can produce well-calibrated *p*-values for spatial co-expression inference, thereby effectively controlling type I error while achieving higher statistical power than the existing methods. We then demonstrated the utility of CONCISE by applying it to multiple ST datasets, including human breast cancer [36], mouse whole embryo [37], mouse model of inflammatory bowel diseases (IBD) [38], and human non-small cell lung cancer (NSCLC) [39] samples profiled using diverse platforms including 10x Visium, Stereo-seq, MERFISH, and CosMx. In analyses of the breast cancer and embryo datasets, where whole-transcriptome profiling enabled the design of evaluation experiments, we showed that spatial autocorrelation and other intrinsic properties of ST data are major confounding factors in spatial co-expression and LRI analyses. By rigorously modeling these factors, CONCISE reduced false-positive discoveries and improved the reliability of inferred co-expression compared with existing approaches. Finally, analyses of mouse IBD model and human NSCLC samples highlight the biological utility of CONCISE in disease contexts. Specifically, CONCISE revealed distinct CCC patterns between inflammation-associated fibroblasts and normal fibroblasts in their interactions with immune cells during intestinal inflammation, and further elucidated signaling interactions sent and received by tumor cells within the NSCLC tumor microenvironment.

## Results

### Method overview

CONCISE infers CCC from context-dependent ST data by identifying L-R pairs whose expression exhibits significant spatially constrained co-expression. To ensure reliable statistical inference, CONCISE introduces a unified measurement-expression modeling framework that accounts for key confounding factors in ST data, including variation in total molecular counts across spots or cells, measurement errors in count observations, and spatial autocorrelation in gene expression (Fig. 1). Explicitly modeling these factors is critical to avoid spurious co-expression signals that may lead to false-positive LRI discoveries.

**Figure 1.**
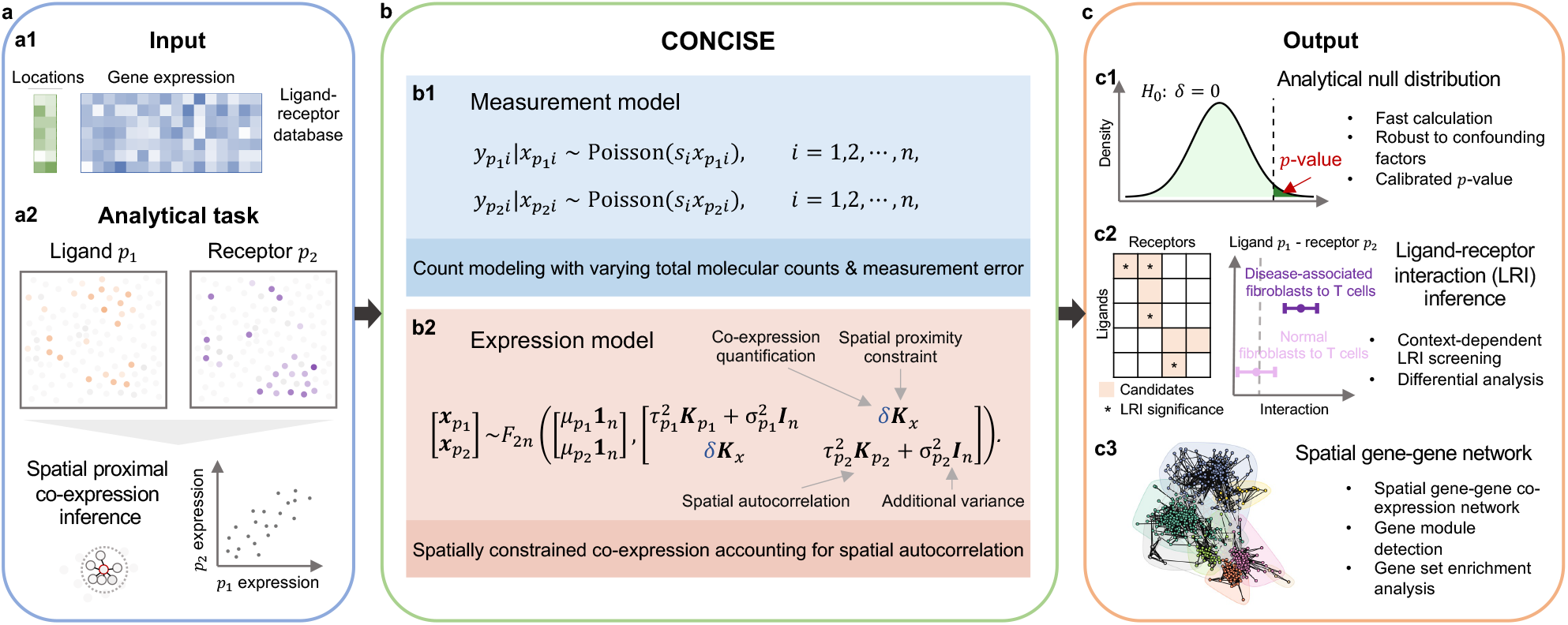
Method overview. CONCISE is a unified statistical framework for spatially proximal co-expression inference and ligand-receptor interaction (LRI) detection from context-dependent ST data. **a.** Workflow of CONCISE. The method takes gene expression count matrix with spatial coordinates as input (**a1**) and identifies LRIs by detecting ligand-receptor co-expression under spatial proximity constraints (**a2**). **b**. Measurement-expression modeling framework. The measurement model links observed counts 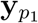 and 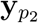 to underlying true expression levels 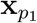 and 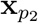, accounting for the count-based nature of ST data, variation in total molecular counts, and measurement errors (**b1**). The latent spatial expression model jointly characterizes ligand and receptor expression, incorporating spatial autocorrelation through spatial kernels 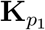 and 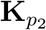, and spatial proximity constraints through **K**_*x*_, when estimating co-expression *δ* (**b2**). **c**. Statistical inference and downstream analyses. A moment-based framework enables efficient parameter estimation and analytical null distribution for fast and well-calibrated *p*-value computation (**c1**). The resulting inference enables LRI screening and comparative analysis of CCC patterns between normal and disease-related cell populations (**c2**), and can also be applied to spatial gene-gene network analysis, supporting spatial gene module identification and gene set enrichment analysis (**c3**).

CONCISE takes as input the observed gene expression count matrix and the spatial coordinates of spots or cells (Fig. 1**a**, panel **a1**). For a candidate L-R pair from a curated database, such as ligand *p*_1_ and receptor *p*_2_, CONCISE models their observed transcript counts 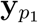and 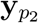 using a measurement model (Fig. 1**b**, panel **b1**) [34, 40]. In this model, latent variables 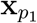 and 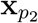 represent the underlying true expression levels of the ligand and receptor, and the observed counts depend on these latent levels as well as the total molecular counts {*s*_*i*_}_*i*_. This formulation captures the count-based nature of ST data, separates the underlying true expression levels and measurement errors in the observed counts, and adjusts for varying total molecular counts across spatial locations.

To quantify spatially constrained co-expression (Fig. 1**a**, panel **a2**), CONCISE further introduces a latent spatial expression model that jointly models 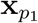 and 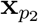 across *n* spatial locations (Fig. 1**b**, panel **b2**). Co-expression is quantified by a parameter *δ*. The model has three key features. First, it accounts for spatial autocorrelation in gene expression by incorporating spatial kernels 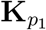 and 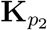 with adaptive scaling factors 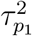 and 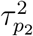. These components model spatial variation in expression that is dependent of the spatial locations. Additional spatially independent variation is modeled by 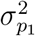 and 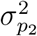 through identity covariance matrices **I**_*n*_. Notably, accounting for spatial autocorrelation is essential for correctly quantifying the uncertainty of co-expression estimates, avoiding inflated false-positive results. Second, the spatial proximity constraints are incorporated through the design of matrix **K**_*x*_. For instance, **K**_*x*_ = **I**_*n*_ corresponds to measuring co-expression within the same spot or cell, analogous to constructing spatial gene-gene networks. For LRI inference, **K**_*x*_ is by default defined by the neighborhood graph, i.e., **K**_*x*_(*i, j*) = 1 only if spots or cells *i* and *j* are spatially proximate, restricting ligand-receptor co-expression inference to nearby spots or cells. Third, the model flexibly characterizes latent expression levels using an unknown nonnegative 2*n*-variate distribution *F*_2*n*_, without restricting a distributional family. For example, if 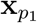 and 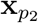 follow Gamma distributions, the resulting measurement-expression model reduces to a negative binomial observation model commonly used for transcript counts. Together, these components form a unified framework for reliable inference of spatially constrained co-expression.

Estimating co-expression and performing statistical inference under a model that accounts for multiple confounding factors is challenging. Without imposing distributional assumptions on *F*_2*n*_, we propose a moment-based framework for efficient parameter estimation and statistically rigorous inference (Fig. 1**c**, panel **c1**). In particular, we derive the analytical null distribution of the test statistic, enabling fast and well-calibrated *p*-value computation for assessing spatially constrained co-expression. The resulting algorithm enables reliable and efficient LRI screening from context-specific ST data, and facilitates detailed comparative analyses of CCC patterns between normal and disease-associated cell populations (Fig. 1**c**, panel **c2**). This flexible framework also accommodates the construction of spatial gene-gene co-expression networks by specifying **K**_*x*_ = **I**_*n*_, enabling downstream analyses such as spatial gene module identification and gene set enrichment analysis (Fig. 1**c**, panel **c3**). Details are included in the Methods section.

### Spatial autocorrelation confounds spatially constrained co-expression analyses and is inherent to spatial transcriptomics data

Spatial autocorrelation is a fundamental concept in spatial statistics, describing the tendency for observations at nearby locations to exhibit similar values. Such spatial dependence arises naturally in spatially resolved data and has been widely recognized across application domains such as geospatial analysis and neuroimaging [24, 27, 29]. If not properly accounted for, spatial autocorrelation can substantially confound downstream statistical analyses, including spatial co-expression inference [26, 27, 28, 29].

To illustrate this effect, we conducted a simulation experiment. For clarity and to avoid introducing additional confounding factors inherent to count data, we generated observations from Gaussian distributions. Specifically, we simulated two independent variables at *n* = 2, 500 spatial locations according to 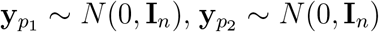. To mimic the spatial dependencies commonly observed in spatially resolved data, we followed established approaches [27, 29] to spatially smooth the random values of 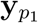 and 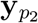 across neighboring locations, generating 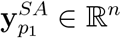 and 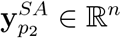, where “SA” denotes spatial autocorrelation. In both settings, 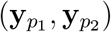 and 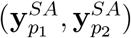, the two variables were independently generated and therefore represent null data for spatial co-expression analysis.

We first examined within-location co-expression analysis, corresponding to the construction of spatial gene-gene networks. For illustration, we compared Pearson’s correlation with the Gaussian variant of CONCISE under the setting **K**_*x*_ = **I**_*n*_. When neither variable exhibits spatial autocorrelation (comparing 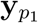 and 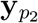), or when only one variable shows spatial dependencies (comparing 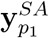 and 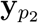), Pearson’s correlation and CONCISE produced consistent co-expression estimates with well-calibrated *p*-values and controlled type I error rates (Fig. 2**a, b**; Supplementary Fig. 1**a, b**). When both variables exhibit spatial autocorrelation (comparing 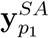 and 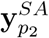), the two methods again produced consistent estimates of co-expression. However, Pearson’s correlation substantially underestimated the uncertainty associated with these estimates relative to CONCISE (Fig. 2**c**). This underestimation leads to spurious co-expression signals and, consequently, inflated false-positive results (Supplementary Fig. 1**c**).

**Figure 2.**
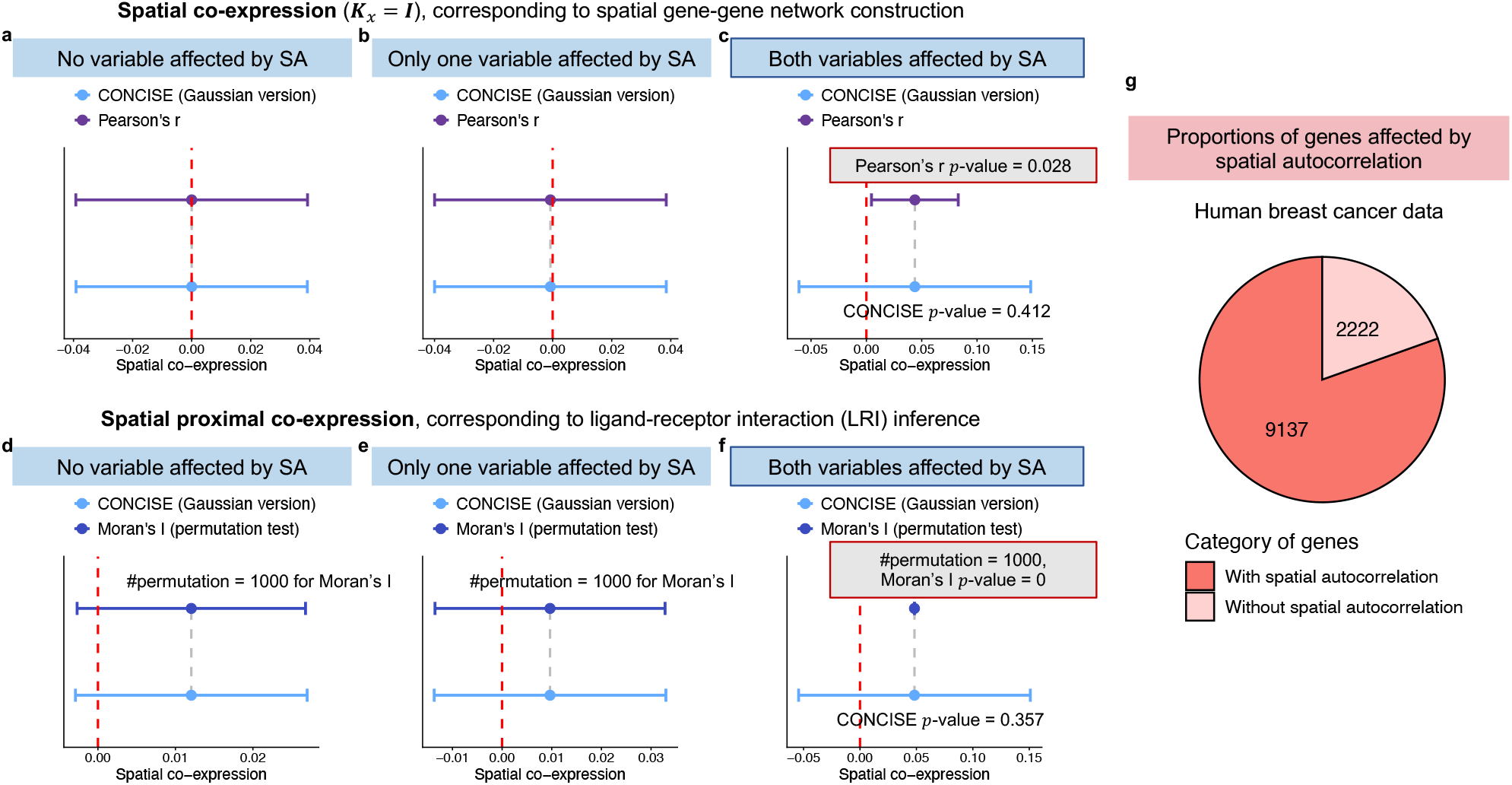
Spatial autocorrelation confounds spatial co-expression and ligand-receptor interaction inference. **a**-**c.** Spatial co-expression analysis under **K**_*x*_ = **I**_*n*_, corresponding to spatial gene-gene network construction. Two independent variables were simulated under three null scenarios: neither variable (**a**), only one variable (**b**), or both variables (**c**) affected by spatial autocorrelation (SA). Pearson’s correlation and the Gaussian version of CONCISE yielded comparable co-expression estimates across all scenarios. When both variables exhibited SA (**c**), Pearson’s correlation substantially underestimated uncertainty and produced spuriously significant results, whereas CONCISE remained well calibrated. **d**-**f**. Spatially proximal co-expression analysis, corresponding to ligand-receptor interaction (LRI) inference. Spatially proximal co-expression was assessed using bivariate Moran’s *I* with permutation testing and CONCISE under the same three null scenarios. When both variables exhibited SA (**f**), the Moran’s *I*-based approach underestimated uncertainty and produced false-positive LRI signals, whereas CONCISE maintained calibrated inference. **g**. Proportion of genes with significant spatial autocorrelation in the human breast cancer Visium dataset [36]. Among 11,359 genes passing sparsity-based quality control, 9,137 genes (80.4%) showed significant spatial autocorrelation.

We next considered spatially proximal co-expression analysis, the setting essential for spatial LRI inference. Many existing methods detect LRIs using statistics derived from bivariate Moran’s *I* [19, 21]. To examine the implications of spatial autocorrelation in this context, we compared a bivariate Moran’s *I*-based approach with CONCISE. In the Moran’s *I* approach, spatially proximal co-expression is quantified using bivariate Moran’s *I* and significance is assessed using permutation-based *p*-values. When at least one of the variables is free from spatial autocorrelation, both approaches produced consistent co-expression estimates and well-calibrated statistical inference. With 1,000 permutations for Moran’s *I*, the two methods also yielded comparable uncertainty quantification (Fig. 2**d, e**; Supplementary Fig. 2**a, b**). However, when both variables exhibit spatial autocorrelation, the Moran’s *I* approach substantially underestimated the uncertainty (Fig. 2**f**). This underestimation again results in spurious signals and inflated false-positive discoveries. In contrast, by explicitly modeling spatial dependencies in the data, CONCISE provided reliable uncertainty quantification and reduced false-positive discoveries (Supplementary Fig. 2**c**).

These results demonstrate that spatial autocorrelation can substantially distort statistical inference in spatial interaction analyses, which underpin many important biological applications. To evaluate the potential impact of this issue in real ST data, we next examined the prevalence of spatial autocorrelation in a human breast cancer dataset profiled by Visium [36] as an example. We quantified the proportion of genes exhibiting spatial autocorrelation in their expression patterns, characterized by significantly non-zero spatial variation in gene expression (i.e., 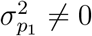 for gene *p*_1_; Fig. 1**b**, panel **b2**). Among the 11,359 genes that pass sparsity-based quality control, more than 80% show significant spatial autocorrelation (Fig. 2**g**). This widespread spatial dependency increases the risk of spurious discoveries when spatial autocorrelation is ignored by existing methods. This observation also highlights the need to model spatial autocorrelation, as in CONCISE, for reliable LRI inference and spatial gene-gene co-expression analysis.

### CONCISE achieves superior false positive rate control in real data permutation studies

To assess false positive rate control in spatial LRI inference, we conducted permutation-based analyses on the breast cancer Visium dataset [36] to generate null datasets in which spatially proximal co-expression is absent. Using these data, we evaluated type I error control of CONCISE and benchmarked its performance against representative state-of-the-art methods, including MERINGUE [18], SpatialDM [19], Copulacci [20], and LIANA+ [21].

To systematically characterize the confounding effects of varying total molecular counts, measurement errors, and spatial autocorrelation on spatial LRI inference, we focused on genes showing spatial autocorrelation (Fig. 2**g**) and designed two experimental scenarios to disentangle these effects (Fig. 3**a**). In Scenario 1, variation in total molecular counts and measurement errors inherent to the data were preserved, while confounding effect from spatial autocorrelation was removed. In Scenario 2, spatial autocorrelation was reintroduced to quantify its additional confounding impact.

**Figure 3.**
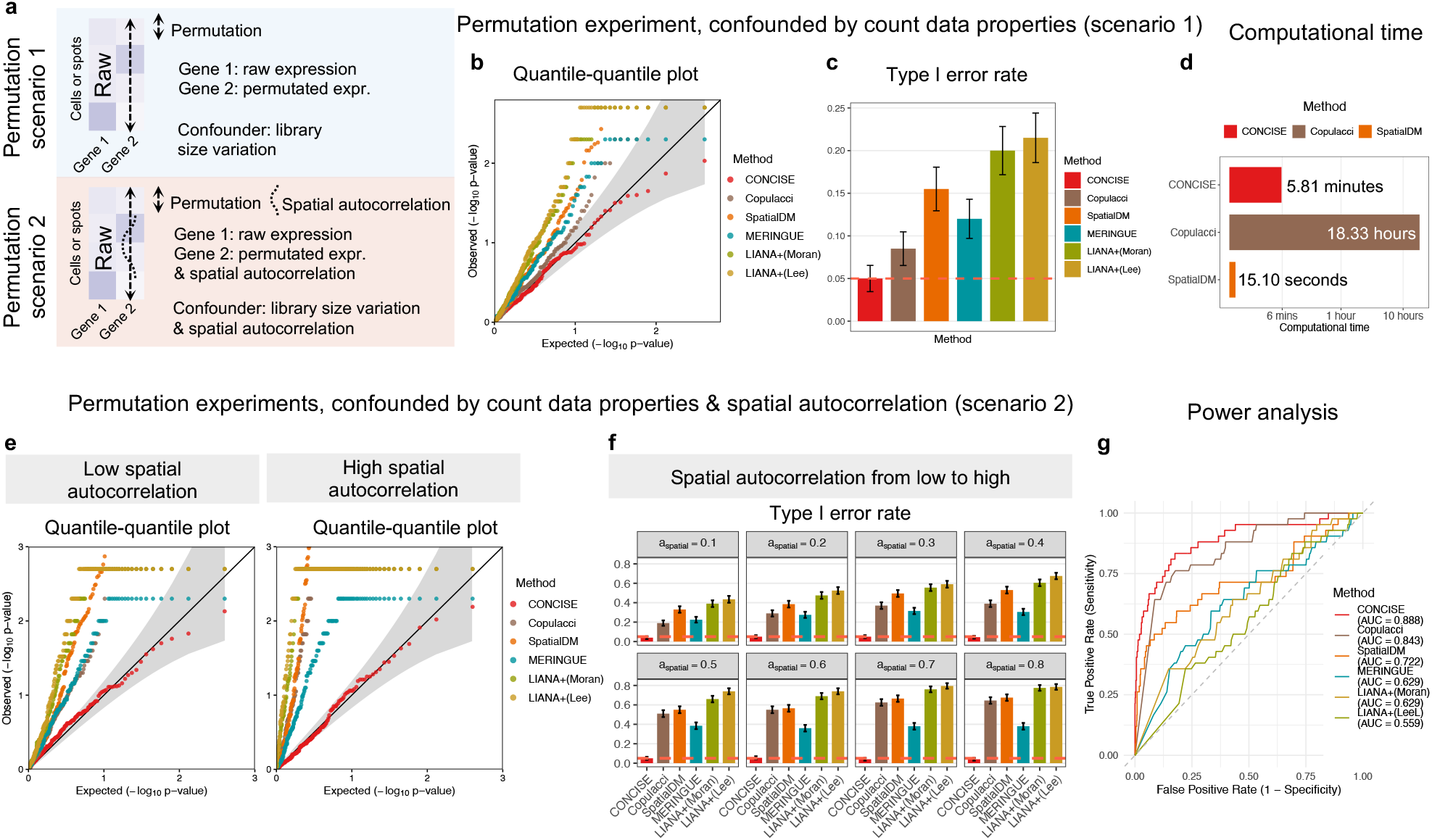
Real data-based permutation study and power analysis. **a.** Schematic overview of two permutation scenarios designed to dissect the multiple confounding effects on spatial LRI inference. Scenario 1 preserves variation in total molecular counts and measurement errors while eliminating confounding from spatial autocorrelation. In Scenario 2, spatial autocorrelation of controlled strength is additionally imposed on the permuted gene, enabling systematic evaluation of its impact on LRI inference. **b**. Quantile-quantile plots of *p*-values from CONCISE and competing methods under Scenario 1. **c**. Comparison of type I error rates across methods under Scenario 1. **d**. Computational time required to analyze 1,000 L-R pairs. **e**. Representative quantile-quantile plots of *p*-values across methods under Scenario 2 at low and high spatial autocorrelation strengths. **f**. Type I error rates under Scenario 2 across increasing levels of spatial autocorrelation. **g**. Power analysis based on simulated null and alternative LRIs parameterized using the Visium breast cancer dataset. Receiver operating characteristic (ROC) curves are shown for all methods, with the area under the ROC curve (AUC) used to quantify inference accuracy.

In Scenario 1, for each randomly sampled gene pair, we retained the raw counts of one gene and permuted the expression of the other across spatial locations to remove spatial co-expression. Specifically, counts were first normalized by total molecular counts at each location and then permuted across locations, after which count data were regenerated using the original total molecular counts. This procedure preserves variation in total molecular counts and measurement errors while eliminating spatial autocorrelation in one gene from each pair. Indicated by the simulation study (Fig. 2; Supplementary Fig. 2), spatial LRI inference under this setting is not confounded by spatial autocorrelation. Under this null setting, CONCISE uniquely achieved well-controlled type I error rates (Fig. 3**c**) and well-calibrated *p*-values (Fig. 3**b**). In contrast, MERINGUE and LIANA+ (bivariate Moran’s *I* and Lee’s *L* variants) operate on normalized or log-normalized counts with permutation-based inference and do not explicitly model measurement errors in the count-generating process. Moreover, their normalization schemes fail to adequately account for variation in total molecular counts, leading to spurious co-expression patterns in both normalized and log-normalized data (Supplementary Fig. 3) [35] and consequently inflated *p*-values and type I error rates. SpatialDM improves computational efficiency and *p*-value estimation by deriving an analytical null distribution under a Gaussian assumption on log-normalized data. However, it remains similarly susceptible and produced inflated type I error. Copulacci, by contrast, adopts a count-based modeling framework. However, it explicitly accounts only for varying total molecular counts, while not effectively modeling measurement errors, resulting in improved yet insufficient false positive control.

We next considered Scenario 2, in which spatial autocorrelation was additionally introduced. Comparison with Scenario 1 isolates the impact of spatial autocorrelation on spatial LRI inference. For each randomly sampled gene pair, we followed the same procedure as in Scenario 1, retaining the raw counts of one gene and permuting the normalized expression of the other across spatial locations. Spatial autocorrelation of controlled strength, parameterized by *a*_spatial_, was then imposed on the permuted expression. Specifically, the mean expression level of each gene was preserved, while spatial variation was set to account for 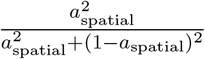 of the total variance. Even when one gene retained its original counts, and only weak spatial autocorrelation (*a*_spatial_ = 0.1) was introduced to the other, all competing methods except CONCISE were substantially confounded, as evidenced by inflated *p*-values and elevated type I error rates relative to Scenario 1 (Fig. 3**e, f**). As the strength of spatial autocorrelation increased, these methods showed progressively higher false positive rates, underscoring the necessity of explicitly modeling spatial autocorrelation in spatial LRI inference. In contrast, CONCISE consistently produced well-calibrated *p*-values and maintained appropriate type I error control across all levels of spatial autocorrelation, highlighting its robust control of false positives in spatial LRI inference.

### CONCISE achieves better detection power

In this section, we evaluated the detection power of spatial LRI inference methods using simulations parameterized from the breast cancer data. Specifically, we generated 2,000 genes with varying mean expression levels and both spatial and non-spatial variance components estimated from the real data. These genes were randomly paired to form 1,000 gene pairs, among which truly spatially constrained co-expressed pairs were defined according to the interactions inferred from the real data. Consistent with the results of the permutation analyses, CONCISE was the only method that maintained proper false positive control, whereas all competing methods exhibited substantially inflated type I error rates (Supplementary Fig. 4). To enable a fair comparison of detection performance under different levels of false positive control, we evaluated receiver operating characteristic (ROC) curves. As shown in Fig. 3**g**, CONCISE achieved the highest area under the curve (AUC), highlighting its superior detection power. Taken together, these results demonstrate that CONCISE uniquely combines rigorous false positive control with high statistical power for spatial LRI inference.

Finally, we compared the computational efficiency of representative methods, benchmarking CONCISE against SpatialDM and Copulacci (Fig. 3**d**). SpatialDM derives an analytical null distribution to enable efficient *p*-value computation, whereas Copulacci adopts a count-based modeling framework and presented comparatively better performance than Gaussian-based approaches. Both CONCISE and SpatialDM completed the analysis of 1,000 gene pairs within six minutes. In contrast, Copulacci required more than 18 hours on four CPU cores, despite using only 200 permutations for *p*-value estimation. Notably, owing to the moment-based estimation framework and explicit derivation of null distributions, CONCISE achieves superior statistical performance while maintaining high computational efficiency.

### Application to Visium breast cancer dataset: negative control study and biological discoveries

In this section, we apply CONCISE to analyze the Visium breast cancer dataset [36]. We first introduce a negative control study based on biological prior knowledge, providing an model-assumption-free benchmark for fairly assessing false positive control across methods. Using this dataset, together with an additional Stereo-seq dataset [37] analyzed below, we demonstrate that CONCISE can consistently achieve desired control of false positives in spatial LRI inference. Next, we focus on spatial gene-gene co-expression analysis and spatial LRI identification in this tumor sample. We show that CONCISE can yield more reliable spatial co-expression detection results than competing methods and is able to identify biologically meaningful LRIs, particularly at tumor boundaries.

We begin with the negative control analysis, which enables a fair evaluation of competing methods directly on the real data. This design is motivated by the fact that expressions of housekeeping genes (HKGs) likely remain stable and are minimally influenced by ligand, receptor activities in neighboring cells [41]. Accordingly, we treat HKGs as negative control receptors and ligands. Under this framework, spatial LRI inference between HKGs and ligands, as well as between HKGs and receptors, is expected to yield no or only a minimal number of significant interactions with appropriate false positive control.

We curated eleven human HKGs reported in a published study [41], including *C1orf43, CHMP2A, EMC7, GPI, PSMB2, PSMB4, RAB7A, REEP5, SNRPD3, VCP*, and *VPS29*. Owing to whole-transcriptome profiling in the Visium dataset, all selected HKGs are present. We applied CONCISE, together with two representative methods, Copulacci and SpatialDM, to infer spatial interactions between these HKGs and a curated set of 69 ligands and 68 receptors from CellChatDB [14] that also passed sparsity-based quality control. The numbers of significant interaction pairs identified by each method are shown in Fig. 4**a, c**. Both Copulacci and SpatialDM reported considerable numbers of significant interactions under this negative control setting. Among 759 HKG-ligand pairs, they identified 64.2% and 66.0% as significant, respectively. Similarly, among 748 HKG-receptor pairs, they reported 58.3% and 67.0% as significant. These findings indicate serious inflation of false positives, consistent with observations from the permutation-based analyses. In contrast, CONCISE identified no or only one significant interaction in these settings, demonstrating reliable control of false positives.

**Figure 4.**
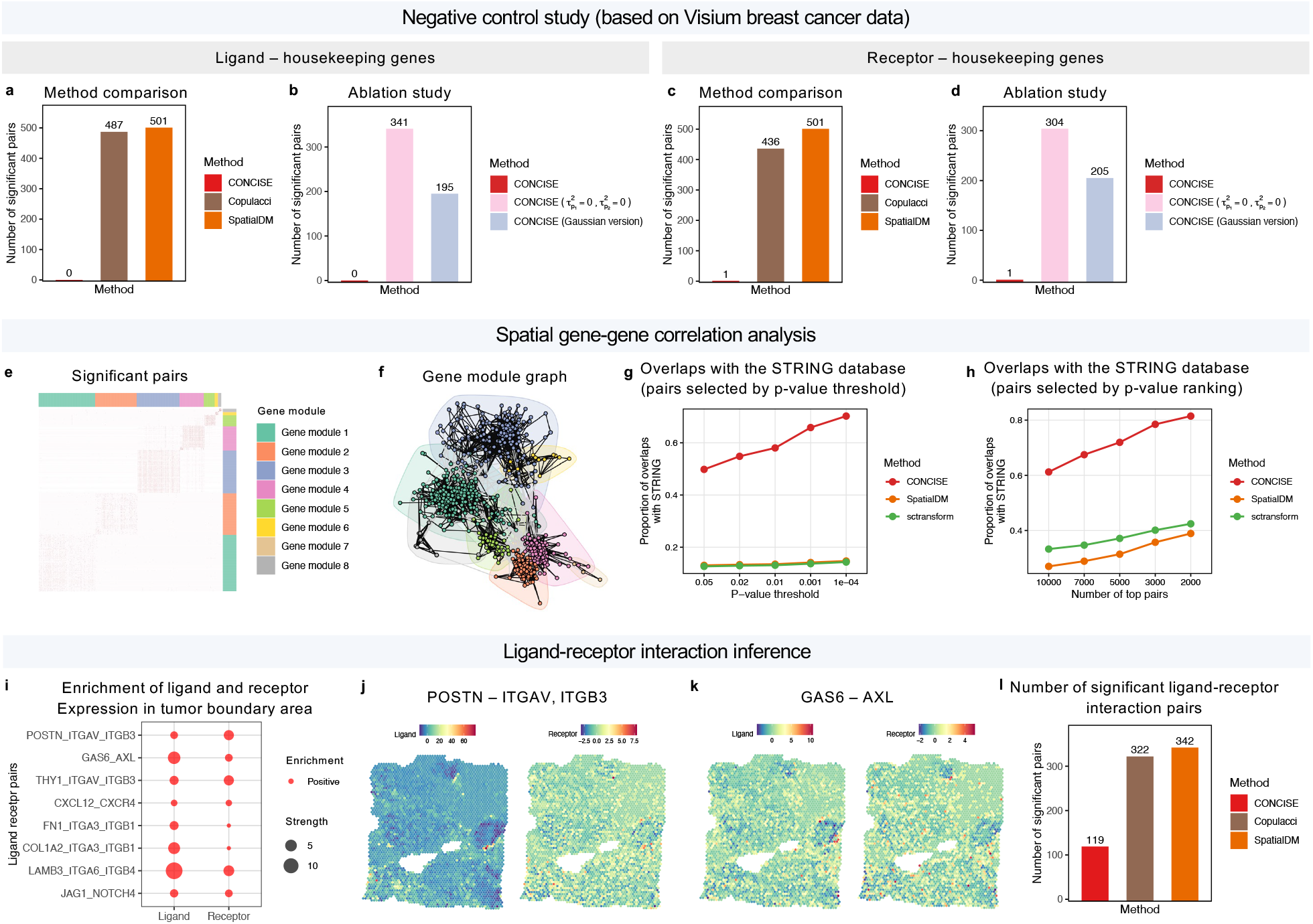
Application of CONCISE to the Visium breast cancer dataset. **a.** Number of significant interactions identified in the ligand-housekeeping gene (HKG) negative control analysis, in which no ture interactions are expected. **b**. Ablation study of CONCISE in the ligand-HKG negative control setting. Spatial variance components were set to zero 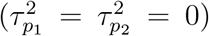 to remove modeling of spatial autocorrelation. The Gaussian variant of CONCISE does not model count data properties. **c**. Number of significant interactions identified in the receptor-HKG negative control analysis, in which no true interactions are expected. **d**. Ablation study of CONCISE in the receptor-HKG negative control setting. **e**. Significant spatial gene-gene co-expression network inferred by CONCISE from the top 1,000 highly variable genes. **f**. Gene modules identified from the inferred spatial gene-gene network. **g**. Proportion of inferred gene pairs overlapping known interactions in the STRING database across different *p*-value significance thresholds, where a higher proportion indicates greater biological relevance. **h**. Proportion of inferred gene pairs overlapping known interactions in the STRING database among the top-ranked gene pairs. **i**. Enrichment analysis of ligand and receptor expression at tumor boundary regions. Representative ligand-receptor pairs preferentially enriched at tumor boundaries are shown. **j**. Spatial expression patterns of the POSTN-ITGAV/ITGB3 interaction. **k**. Spatial expression patterns of the GAS6-AXL interaction. **l**. Number of significant spatial LRIs identified by different methods.

To elucidate the mechanisms underlying this robustness, we conducted an ablation study of CONCISE. First, we manually set the spatial variance components to zero, i.e., 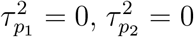 yielding a variant of CONCISE that does not account for spatial autocorrelation. This ablated model identified over 300 significant pairs in both HKG-ligand and HKG-receptor negative control settings (Fig. 4**b, d**), demonstrating severe inflation of false positives when spatial autocorrelation is ignored. We next applied the Gaussian variant of CONCISE that does not model the count nature of the data, variation in total molecular counts, and measurement errors. This variant identified approximately 200 significant pairs in the same negative control settings, highlighting the additional confounding effects of varying total molecular counts and measurement errors. Together, these results establish spatial autocorrelation, variation in total molecular counts, and measurement errors as major sources of confounding in real data. By explicitly modeling all three, CONCISE uniquely achieves well-calibrated inference and effective control of false positives.

Next, we evaluated the biological credibility of the significant relationships identified by CONCISE. Direct validation of inferred CCC events is challenging because comprehensive ground-truth datasets are largely unavailable. Instead, we considered a special case in which the interaction kernel was set to **K**_*x*_ = **I**. Under this setting, CONCISE reduces to a framework for inferring spatial gene-gene co-expression within individual cells or spots, allowing the inferred relationships to be evaluated against known biological interaction networks curated in STRING [42], where a higher overlap indicates greater detection reliability and biological relevance. Specifically, we inferred spatial co-expression relationships among the top 1,000 variable genes in the dataset, yielding 499,500 gene pairs in total. Given the heavy computational burden of Copulacci for analyzing these 499,500 gene pairs, we excluded it from this comparison. In addition to SpatialDM, we included the sctransform-based approach [33] for a more comprehensive evaluation. Specifically, sctransform was used to estimate expression levels from count data by correcting for total molecular counts under a negative binomial model, followed by Pearson’s correlation on the resulting residuals to infer gene-gene co-expression. This approach accounts for key properties of count data but does not model spatial autocorrelation. Across a wide range of *p*-value thresholds, CONCISE consistently achieved substantially higher overlap with the STRING database than competing methods (Fig. 4**g**), indicating better detection reliability. To account for differences in the number of significant pairs identified by different methods, we further performed a ranking-based comparison. Gene pairs were ranked by significance, and the top-ranked pairs (ranging from 2,000 to 10,000) were evaluated. Consistent with the threshold-based analysis, CONCISE achieved consistently higher overlap with STRING across all settings (Fig. 4**h**), further demonstrating its better ability to identify truly interacting gene pairs.

After validating the better performance of CONCISE, we next analyzed the 15,826 significant gene pairs identified at a multiple-testing-adjusted *p*-value threshold of 0.05. These pairs define a spatial gene-gene network, shown in Fig. 4**e**. Applying community detection algorithm [43] to this network revealed eight distinct gene modules (Fig. 4**f**). We found four modules (modules 1, 2, 4, and 6) spatially localized near the tumor regions with different spatial patterns (Supplementary Fig. 5). Gene ontology (GO) analysis indicated that they capture key biological processes associated with tumor microenvironments (Supplementary Fig. 6). Specifically, modules 1 and 4 were enriched for hypoxia response and blood vessel development, respectively, consistent with hypoxic conditions and angiogenic activity at tumor margins [44, 45, 46]. In addition, modules 2 and 6 were enriched for immune-related processes, including positive regulation of immune response, T cell activation, granulocyte chemotaxis, and leukocyte migration involved in inflammatory response. The presence of these modules indicate active immune cell recruitment and infiltration at the tumor-stroma interface [47, 48], supporting the high quality and biological interpretability of CONCISE.

Having established the reliability of CONCISE through multiple validation experiments, we finally applied it to infer LRIs from the human breast cancer dataset. Among the curated set of 612 candidate L-R pairs from CellChatDB that passed sparsity-based quality control, CONCISE identified 119 pairs exhibiting significant spatial interaction signals after multiple testing correction (Fig. 4**l**). By contrast, in negative control experiments, CONCISE detected nearly no interactions across two control sets, each comprising over 700 pairs. The enrichment of discoveries among curated L-R pairs relative to negative controls provides additional evidence that CONCISE can reliably distinguish biologically meaningful CCC signals from spurious spatial correlations. The identified interactions revealed biologically meaningful insights into tumor biology. In particular, we identified tumor boundary regions based on the pathological annotation (Supplementary Fig. 7) and investigated LRIs preferentially localized to these interfaces through enrichment analysis. Representative L-R pairs are shown in Fig. 4**i**. Among them, we observed enrichment of the CXCL12-CXCR4 signaling axis near tumor boundaries, consistent with its established roles in tumor cell proliferation, angiogenesis, and immune evasion [49, 50]. We also identified enrichment of GAS6-AXL signaling (Fig. 4**k**), a pathway increasingly recognized as an essential mediator of tumor-cell invasion and metastatic progression [51, 52]. Inhibition of the GAS6/AXL axis is known to be critical to suppress tumor growth across cancers, and AXL itself has emerged as a promising therapeutic target for overcoming tumor progression and therapeutic resistance [53, 54]. In addition, POSTN-ITGAV/ITGB3 signaling showed activity at tumor boundaries, consistent with the roles of periostin and integrin *α*V*β*3 in stromal remodeling, angiogenesis, and metastasis [55, 56]. In summary, these findings support that CONCISE can identify spatial LRIs with strong biological relevance and uncover key signaling programs associated with tumor progression and microenvironmental remodeling.

### Application to Stereo-seq mouse embryo dataset: negative control study and biological discoveries

To further evaluate the robustness and generalizability of CONCISE in a distinct biological context, we applied it to a high-resolution Stereo-seq mouse whole-embryo dataset [37]. Following the analysis of the Visium human breast cancer dataset, we first performed a negative control study based on HKGs to assess false positive control directly on the real embryo data. We then leveraged the flexibility of CONCISE to conduct spatial gene-gene co-expression analysis, where the reliability of the identified relationships was evaluated through their overlap with known biological network in the STRING database. Finally, we applied CONCISE to infer spatial LRIs and characterize developmental communication pathways across the embryo.

We adopted the same negative control strategy used in the human breast cancer analysis. For the mouse embryo dataset, we curated 12 well-established mouse HKGs [57], including *Actb, Atp5f1, B2m, Gapdh, Hprt, Pgk1, Rer1, Rpl13a, Rpl27, Sdha, Tbp*, and *Ubc*. All of these genes were captured by the Stereo-seq platform owing to its whole-transcriptome profiling capability. Treating these HKGs as negative-control receptors and ligands, we evaluated spatial interactions between them and curated ligand or receptor genes using CONCISE, Copulacci, and SpatialDM. As HKG expression is expected to be minimally affected by ligand and receptor activities in neighboring cells, significant interactions identified under this design are likely to represent false positive discoveries. Consistent with the findings from the human breast cancer data, Copulacci and SpatialDM identified a substantial number of significant interactions among the 360 HKG-ligand pairs and 396 HKG-receptor pairs, declaring up to 45.5% and 36.1% of these pairs significant, respectively (Fig. 5**a, c**). These results indicate serious inflation of false positive findings. In contrast, CONCISE detected fewer than ten significant interactions across all tested pairs, demonstrating improved false positive control.

**Figure 5.**
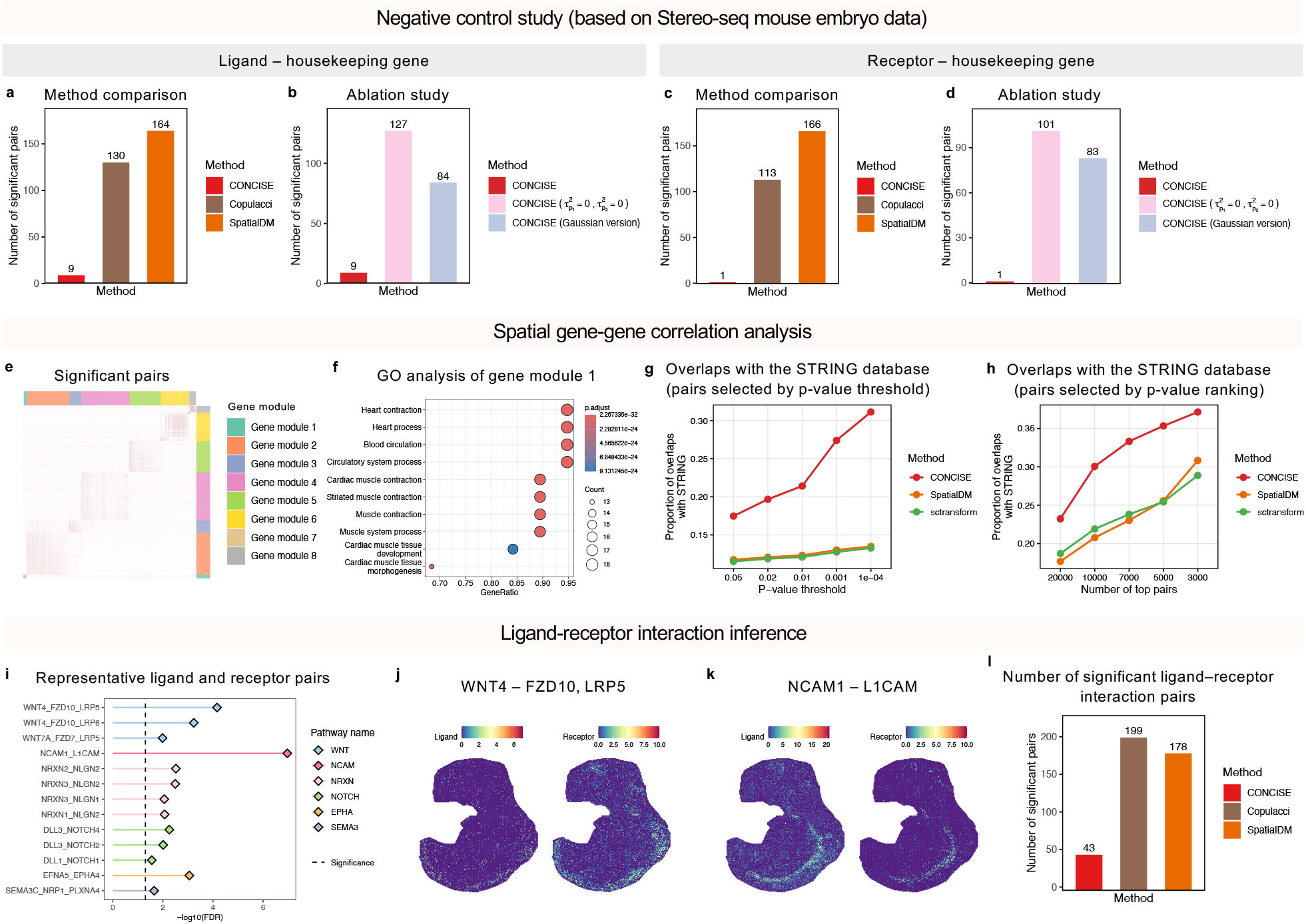
Application of CONCISE to the Stereo-seq mouse embryo dataset. **a.** Number of significant interactions identified in the ligand-HKG negative control analysis, in which no true interactions are expected. **b**. Ablation study of CONCISE in the ligand-HKG negative control setting. **c**. Number of significant interactions identified in the receptor-HKG negative control analysis. **d**. Ablation study of CONCISE in the receptor-HKG negative control setting. **e**. Significant spatial gene-gene co-expression network inferred by CONCISE from the top 1,000 highly variable genes. **f**. Gene ontology (GO) enrichment analysis of gene module 1, revealing enrichment for cardiac development and function. **g**. Proportion of inferred gene pairs overlapping known interactions in the STRING database across different significance thresholds, where a higher proportion indicates greater biological relevance. **h**. Proportion of inferred gene pairs overlapping known interactions in the STRING database among the top-ranked gene pairs. **i**. Representative significant LRIs identified by CONCISE. Interactions are grouped according to their associated signaling pathways. **j**. Spatial expression patterns of the WNT4-FZD10/LRP5 interaction. **k**. Spatial expression patterns of the NCAM1-L1CAM interaction. **l**. Number of significant spatial LRIs identified by different methods.

To further investigate the source of this improvement, we repeated the ablation analysis of CONCISE on the Stereo-seq mouse embryo dataset. The results corroborated those observed in the human breast cancer dataset and further supported the contribution of the methodological innovations underlying CONCISE’s improved false positive control. Specifically, when the component accounting for spatial autocorrelation was removed by setting the spatial variance components to zero 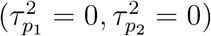, the resulting variant of CONCISE identified more than 100 significant pairs in both the HKG-ligand and HKG-receptor negative control analyses, corresponding to approximately 35% and 25% of the tested pairs, respectively. Similarly, the Gaussian variant of CONCISE, which does not account for the count-based nature of ST data, heterogeneous total molecular counts across spatial locations, and measurement errors, also produced a considerable number of significant findings, including more than 80 pairs in each negative control setting. Collectively, these results again demonstrate the necessity and effectiveness of the key modeling components introduced in CONCISE for improving the reliability of spatial LRI inference.

We next leveraged the flexibility of CONCISE to investigate spatial gene-gene co-expression patterns during embryonic development. Similar to the breast cancer analysis, we inferred spatial co-expression relationships among the top 1,000 highly variable genes, yielding a spatial gene-gene network that revealed multiple co-expression modules (Fig. 5**e**). These modules corresponded to key developmental processes. For example, gene module 1 was associated with cardiac development and function, whereas gene modules 2 and 3 captured neuronal morphogenesis and embryonic skeletal system development, respectively (Fig. 5**f**; Supplementary Fig. 8). Importantly, this analysis also enabled a quantitative assessment of the biological relevance of the inferred spatial relationships (Fig. 5**g, h**). We evaluated the identified co-expression against known biological networks in the STRING database, where greater overlap indicates higher biological relevance. We compared CONCISE with SpatialDM and an sctransform-based approach, while excluding Copulacci because of its substantial computational burden. Specifically, Copulacci required approximately 10 minutes to analyze a single gene pair using eight CPU cores in the HKG-based analysis, making a comprehensive analysis of all 499,500 gene pairs computationally prohibitive. Across a broad range of *p*-value thresholds, CONCISE consistently achieved higher overlap with STRING than competing methods (Fig. 5**g**). Similar conclusions were obtained from a ranking-based comparison using top-ranked gene pairs, which accounts for differences in the numbers of significant pairs identified by different methods (Fig. 5**h**). Together, these results demonstrate that CONCISE preferentially identifies biologically meaningful spatial gene-gene relationships.

Finally, we applied CONCISE to infer LRIs from the Stereo-seq mouse embryo dataset. Among the 424 curated candidate L-R pairs from CellChatDB that passed sparsity-based quality control, CONCISE identified 43 significant spatial interaction pairs after multiple-testing correction (Fig. 5**l**). In contrast, across two negative-control sets comprising a total of 756 curated pairs, CONCISE detected no more than ten significant interactions. This clear separation between biologically plausible and negative-control pairs further supports the reliability of CONCISE for identifying spatial CCC. The significant LRIs spanned multiple signaling pathways, with representative examples shown in Fig. 5**i**. For example, the WNT4-FZD10/LRP5 interaction was enriched in the developing dorsal neural tube, where its spatial distribution closely overlapped with the expression of regional marker genes, including *Pax3, Pax7*, and *Atoh1* [58, 59, 60] (Fig. 5**j**; Supplementary Fig. 9). These observations are consistent with the established role of Wnt signaling in dorsal neural tube patterning and neuronal differentiation [61, 62, 63]. As another example, NCAM1-L1CAM and NRXN3-NLGN1 interactions were enriched in regions undergoing neuronal development (Fig. 5**k**; Supplementary Fig. 10). These interactions spatially overlapped with the expression of neuronal markers (*Tubb3, Elavl3*, and *Stmn2*) [64, 65, 66], as well as genes involved in synaptic function (*Syt1* and *Snap25*) [67, 68] (Supplementary Fig. 11), suggesting that these communication programs may contribute to neuronal maturation and the establishment of early neural connectivity [69, 70, 71]. These findings further demonstrate the ability of CONCISE to recover biologically meaningful and fine-grained CCC programs associated with embryonic development.

### CONCISE reveals distinct CCC patterns between inflammation-associated cells and normal cells in MERFISH intestinal inflammation section

To further demonstrate the ability of CONCISE to uncover disease-associated CCC, we applied it to a MERFISH section from a dextran-sodium-sulfate (DSS)-induced mouse colitis model at peak inflammation [38]. This mouse model recapitulates key features of inflammatory bowel disease (IBD), a chronic relapsing disorder that affects millions of individuals worldwide and remains an unmet medical challenge [72, 73]. By capturing the cellular composition and spatial organization of the inflamed gut, it provides an opportunity to investigate how CCC shapes inflammatory microenvironment and tissue remodeling. Understanding these communication programs may provide insights into IBD pathogenesis and identify potential therapeutic targets.

Leveraging the single-cell resolution of MERFISH, the dataset profiles the expression of 940 genes across more than 23,000 cells. It encompasses diverse epithelial, immune, stromal and endothelial cell types, including a distinct population of inflammation-associated fibroblasts (IAFs) characterized by elevated expression of *Egr1, Fos* and *Igfbp5* (Fig. 6**a, e**). Given the central role of fibroblast-immune interactions in intestinal inflammation, we next investigated whether IAFs exhibit communication patterns distinct from those of homeostatic fibroblasts. Notably, CONCISE naturally accommodates cellular-resolution ST data and, when cell-population annotations are available, enables cell-population-specific inference through appropriate specification of the interaction kernel **K**_*x*_. Furthermore, the interaction estimates with calibrated uncertainty quantification produced by CONCISE enable direct statistical comparisons of LRIs across related cell populations within the same biological context. We therefore applied CONCISE to infer spatial LRIs between fibroblasts and immune cells, including T cells, B cells and macrophages, and compared the resulting interaction profiles among IAFs and two major homeostatic fibroblast populations (fibroblast 1 and fibroblast 2). CONCISE revealed a clear enrichment of significant LRIs between IAFs and immune cells relative to homeostatic fibroblasts, a pattern that was consistently observed across T cells, B cells and macrophages (Fig. 6**b-d**; Supplementary Fig. 12). Among T cells, IAF-associated interactions include the well-established chemokine signaling axes CCL8-CCR2 [74] and CXCL12-CXCR4 [75], the TNF-family interaction TNFSF14-TNFRSF14 [76], and the inflammatory cytokine axis IL1B-IL1R1/IL1RAP [77], whereas these interactions were not detected or were markedly weaker for homeostatic fibroblasts (Fig. 6**f**). Similar IAF-enriched patterns were observed for B cells and macrophages, including CXCL12-CXCR4 and TNFSF13B-TNFRSF13B [78] in B cells, and CCL8-CCR2, INHBB-ACVR2B [79] and FGF2-FGFR1 [80] in macrophages (Fig. 6**g,h**). These findings highlight distinct communication programs associated with the inflammatory fibroblast state and suggest that IAFs may act as a signaling hub within the inflamed gut, engaging in enhanced communication with multiple immune populations and potentially contributing to immune-cell recruitment, activation and maintenance during intestinal inflammation.

**Figure 6.**
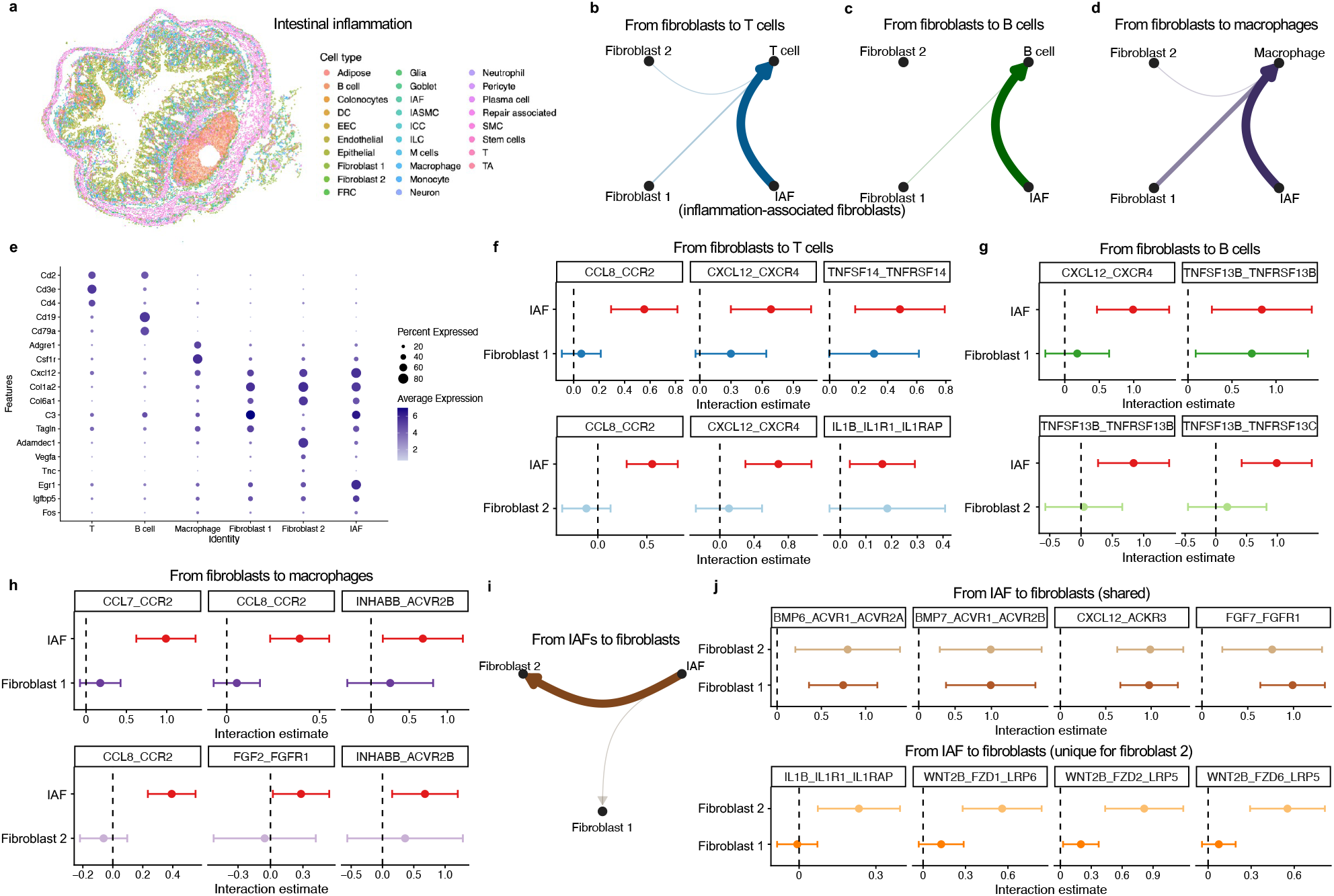
CONCISE reveals distinct spatial LRIs associated with inflammation-associated fibroblasts in DSS-induced colitis. **a.** The MERFISH intestinal inflammation section with cell-type annotation. **b**-**d**. Cell-population-level communication networks from fibroblasts to T cells (**b**), B cells (**c**), and macrophages (**d**). Edge width is proportional to the number of significant LRIs. **e**. Expression of representative marker genes across immune and fibroblast populations. **f**-**h**. Representative LRIs from fibroblasts to T cells (**f**), B cells (**g**), and macrophages (**h**). Interaction estimates inferred by CONCISE are shown for inflammation-associated fibroblasts (IAFs) and homeostatic fibroblast populations. **i**. Cell-population-level communication network from IAFs to homeostatic fibroblast populations. Edge width is proportional to the number of significant LRIs. **j**. Representative LRIs from IAFs to fibroblast populations. Shared interactions detected in both fibroblast populations are shown in the top panel, whereas fibroblast-2-specific interactions are shown in the bottom panel.

Beyond their interactions with immune cells, we next investigated whether IAFs also communicate with other fibroblast populations and thereby contribute to remodeling of the stromal compartment itself. To address this question, we applied CONCISE to infer LRIs from IAFs to the two major homeostatic fibroblast populations. Although IAFs communicated with both fibroblast populations, the interaction landscape was more extensive for fibroblast 2 than for fibroblast 1 (Fig. 6**i**; Supplementary Fig. 13), suggesting that IAF-derived signaling is preferentially directed toward a specific stromal state. Several signaling interactions were shared between fibroblast 1 and fibroblast 2, including CXCL12-ACKR3 [81], FGF7-FGFR1 [80], BMP7-ACVR1/ACVR2B, BMP6-ACVR1/ACVR2A [82] (Fig. 6**j**), indicating the presence of common stromal communication patterns between IAFs and homeostatic fibroblasts. Notably, although fibroblast 2 showed weaker interactions with immune cells, it exhibited an additional set of IAF-derived interactions that were not detected in fibroblast 1, including the inflammatory cytokine axis IL1B-IL1R1/IL1RAP [77] and multiple WNT2B-mediated signaling pathways, such as WNT2B-FZD1/LRP6, WNT2B-FZD2/LRP5 and WNT2B-FZD6/LRP5 [83] (Fig. 6**b**-**d, j**). These fibroblast-2-specific interactions suggest enhanced responsiveness to inflammation-associated signals originating from IAFs. Interestingly, fibroblast 2 show elevated expression of *Adamdec1* and *Vegfa* (Fig. 6**a, e**), genes that have been implicated in tissue remodeling [84, 85]. The preferential communication between IAFs and fibroblast 2 therefore suggests that inflammatory fibroblasts may selectively engage stromal populations associated with remodeling of the inflamed intestinal microenvironment. These findings together indicate that IAFs can serve as a central signaling hub within the inflamed intestine, coordinating communication across both immune and stromal compartments and potentially shaping the local inflammatory microenvironment.

### CONCISE elucidates signaling interactions in NSCLC tumor microenvironment based on the CosMx section

In this section, we further demonstrate the utility of CONCISE for characterizing disease-associated CCC in complex tissues by applying it to a large-scale human non-small cell lung cancer (NSCLC) dataset generated using the CosMx platform [86]. This dataset profiles the expression of 960 genes across 81,236 cells at single-cell resolution. Examination of the spatial organization of the tissue revealed extensive co-localization of tumor cells with diverse immune and stromal populations, including T cells, macrophages, fibroblasts, endothelial cells and mast cells (Fig. 7**a, b**), suggesting abundant opportunities for intercellular communication within the tumor microenvironment.

We applied CONCISE to systematically characterize signaling interactions between tumor cells and their surrounding cell populations. Among interactions originating from tumor cells, CONCISE identified multiple established spatial LRIs involved in tumor progression and immune regulation, with representative examples shown in Fig. 7**e**-**i**. Notably, tumor-to-T-cell communication was marked by CD274-PDCD1 signaling (Fig. 7**e**), corresponding to the canonical PD-1/PD-L1 immune checkpoint pathway that suppresses anti-tumor T-cell responses [87, 88]. Additional interactions, including CDH1-ITGAE/ITGB7 and ICAM1-ITGAL/ITGB2, highlighted direct communication between tumor cells and T cells through pathways involved in T-cell retention, cell adhesion and immune synapse formation [89, 90, 91]. Tumor-to-macrophage communication was dominated by MIF-mediated signaling, including MIF-CD74/CXCR4, MIF-CD74/CD44 and MIF-CD74/CXCR2 interactions, together with VEGFA-VEGFR1 signaling (Fig. 7**f**), pathways known to regulate macrophage recruitment, activation and tumor-supportive immune functions [92, 93, 94]. In addition, tumor cells communicated extensively with fibroblasts through growth factor and inflammatory signaling pathways, including PDGFA-PDGFRA, PDGFC-PDGFRA, IL1A-IL1R1/IL1RAP and FGF2-FGFR1 (Fig. 7**g**), consistent with activation of stromal programs associated with tissue remodeling and tumor progression [95, 96, 97, 98]. Tumor-to-endothelial communication was characterized by VEGFA-VEGFR2 and VEGFA-VEGFR1R2 signaling [94, 99, 100] (Fig. 7**h**), highlighting active angiogenic signaling within the tumor microenvironment, whereas KITL-KIT signaling suggested that tumor cells may also promote the maintenance of mast cells [101, 102] (Fig. 7**i**).

**Figure 7.**
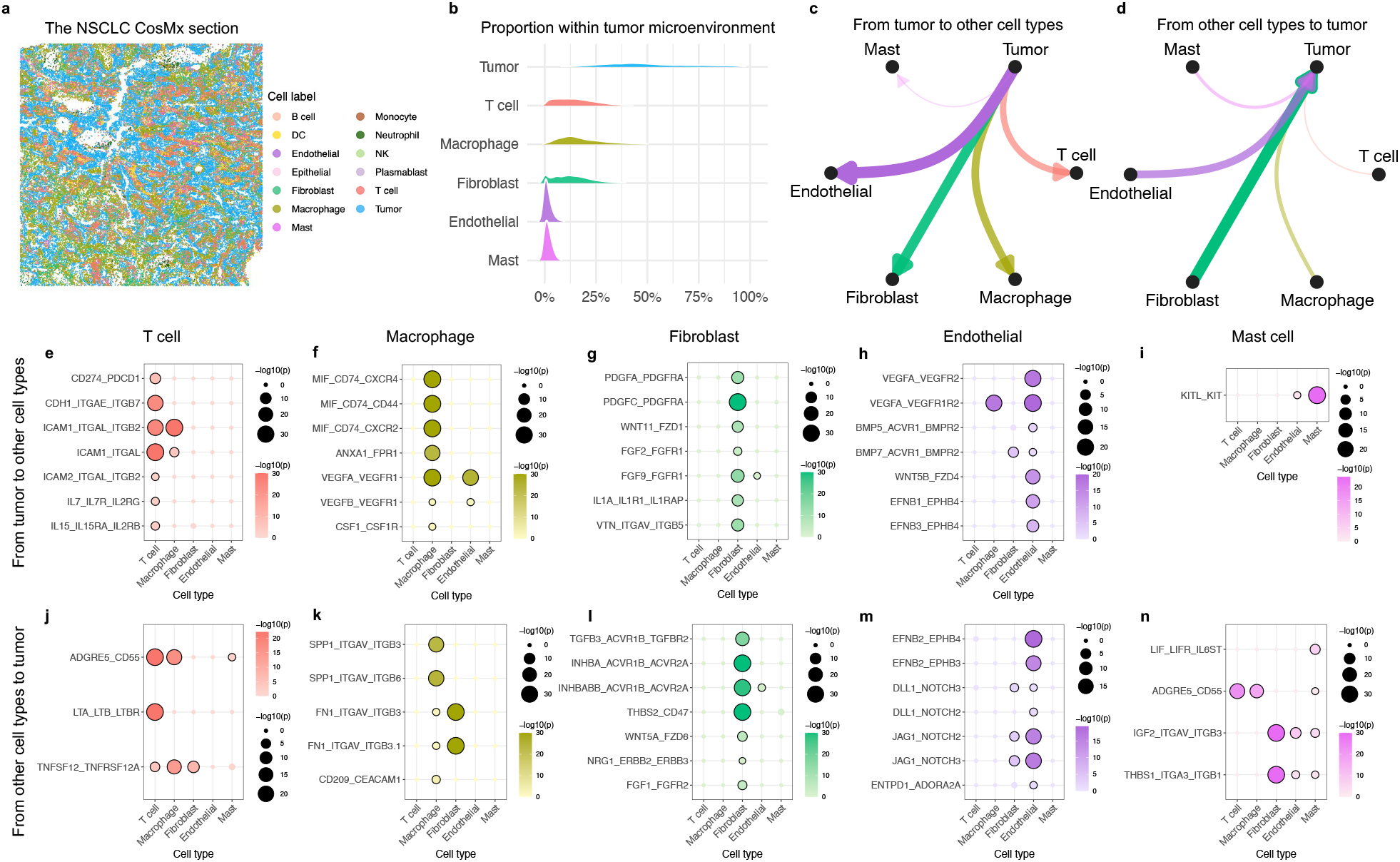
CONCISE elucidates spatial LRIs within the NSCLC tumor microenvironment. **a.** The CosMx NSCLC section with cell-type annotation. **b**. Cell-type composition of the tumor microenvironment. **c**. Cell-population-level communication network summarizing significant LRIs originating from tumor cells. **d**. Cell-population-level communication network summarizing significant LRIs directed toward tumor cells. Edge width is proportional to the number of significant LRIs. **e**-**i**. Representative significant LRIs from tumor cells to T cells (**e**), macrophages (**f**), fibroblasts (**g**), endothelial cells (**h**), and mast cells (**i**). Dot size and color indicate statistical significance. **j**-**n**. Representative significant LRIs from T cells (**j**), macrophages (**k**), fibroblasts (**l**), endothelial cells (**m**), and mast cells (**n**) to tumor cells.

We next examined signaling directed toward tumor cells from surrounding cellular populations. CONCISE identified multiple biologically established pathways through which the tumor microenvironment may influence malignant cells. Macrophage-to-tumor communication was characterized by SPP1- and FN1-mediated integrin signaling, including SPP-ITGAV/ITGB3 and FN1-ITGAV/ITGB3 interactions (Fig. 7**k**), which have been implicated in extracellular matrix remodeling, tumor invasion and malignant progression [103, 104]. Fibroblasts communicated with tumor cells through TGFB3-ACVR1B/TGFBR2, INHBA-ACVR1B/ACVR2A and NRG1-ERBB2/ERBB3 signaling (Fig. 7**l**), highlighting stromal regulation of tumor growth, survival and progression [105, 106, 107]. Endothelial-to-tumor communication was enriched for JAG1-NOTCH2/3, DLL1-NOTCH2/3 and EFNB2-EPHB4 signaling (Fig. 7**m**), consistent with established roles of vascular niche signaling in regulating tumor-cell state, angiogenesis and tumor-vascular interactions [108, 109, 110, 111]. Additionally, CONCISE identified signaling pathways including LTA-LTB/LTBR, TNFSF12-TNFRSF12A, and LIF-LIFR/IL6ST interactions (Fig. 7**j, n**), suggesting direct immune regulation of tumor cells [112, 113, 114].

The recovery of multiple well-established signaling pathways spanning immune regulation, stromal activation, angiogenesis and tumor progression supports the biological validity of the interactions identified by CONCISE. To obtain a global view of intercellular communication within the NSCLC microenvironment, we next summarized the number of inferred spatial LRIs into a cell-population-level communication network (Fig. 7**c, d**). The resulting network revealed distinct directional communication patterns between tumor cells and their surrounding microenvironment. Tumor cells exhibited extensive outgoing signaling toward immune populations, particularly T cells, whereas signaling from T cells back to tumor cells was comparatively limited (Supplementary Fig. 14). In contrast, communication between tumor cells and fibroblasts displayed the opposite trend, with fibroblasts contributing more signaling interactions toward tumor cells than they received. These asymmetric communication patterns suggest different functional roles for immune and stromal interactions within the NSCLC microenvironment. Tumor cells appear to actively shape local immune states and potentially attenuate anti-tumor immune responses, whereas fibroblasts may serve as a major stromal niche that supports tumor progression through extensive signaling directed toward malignant cells.

## Discussion

In this paper, we presented CONCISE, a statistically principled framework for spatially constrained co-expression and LRI inference from spatial transcriptomics data. By jointly accounting for spatial autocorrelation, heterogeneity in total molecular counts, measurement errors, and spatial proximity constraints within a unified modeling framework, while avoiding restrictive assumptions on the underlying expression distributions, CONCISE enables robust and rigorous inference of spatial CCC. Through an analytically derived null distribution, CONCISE further supports computationally efficient statistical testing without relying on computationally intensive permutation procedures. Across extensive simulations, benchmarking analyses, and applications to diverse spatial transcriptomics datasets, we showed that CONCISE achieves well-calibrated statistical inference, effective control of false-positive rates, and enhanced power for detecting biologically meaningful interactions.

Importantly, our study highlights several fundamental challenges underlying spatial CCC inference. Real-data permutation experiments and biologically motivated negative-control analyses across spatial transcriptomics platforms consistently revealed that spatial autocorrelation, variation in total molecular counts, and measurement errors can induce substantial spurious spatial co-expression signals when not properly modeled. Despite their importance, these sources of confounding remain insufficiently addressed by most existing approaches. While many methods incorporate spatial information through neighborhood definitions or distance-based constraints, very few model the spatial dependence of gene expression itself. Furthermore, most approaches operate on normalized expression values and largely ignore the count-generating process underlying observed transcript counts. As a consequence, they often underestimate uncertainty, produce inflated false-positive results, and yield less reliable inference. In contrast, CONCISE addresses all these challenges and achieves substantial improvements in inference reliability while maintaining computational efficiency through key methodological innovations.

The first major advantage of CONCISE lies in its explicit modeling of ligand and receptor expression as spatial processes. Spatial autocorrelation, the tendency for nearby locations to exhibit similar expression, is a pervasive characteristic of spatial transcriptomics data and a fundamental concept in spatial statistics. Research in spatial statistics has established that failure to account for spatial dependence can substantially underestimate uncertainty, leading to spurious false-positive results. Nevertheless, existing spatial CCC methods generally neglect the spatial dependence intrinsic to ligand and receptor expression. By uniquely modeling spatial autocorrelation within a rigorous statistical framework, CONCISE integrates well-established principles from spatial statistics into the analysis of spatially resolved cellular communication, enabling accurate uncertainty quantification, appropriate control of false-positive discoveries, and better identification of biologically meaningful spatial interactions.

Second, CONCISE adopts a measurement-expression modeling framework that explicitly separates latent true expression from observed transcript counts. Rather than treating normalized expression values as direct measurements for gene activity, CONCISE models the underlying count-generating process and accounts for both variation in total molecular counts and measurement uncertainty. Similar considerations have become important in single-cell transcriptomics studies, where variation in total molecular counts and measurement errors are now recognized as major sources of confounding that can substantially affect downstream analyses. Our results demonstrate that these factors likewise distort spatial co-expression and LRI inference. By explicitly modeling the measurement process, CONCISE substantially reduces spurious interaction signals, thereby enabling more reliable inference of spatial CCC.

Finally, CONCISE combines a flexible moment-based estimation framework with analytical hypothesis testing. Unlike existing methods that rely on restrictive distributional assumptions or computationally intensive permutation procedures, CONCISE accommodates a broad class of latent expression distributions while retaining analytical tractability. This formulation enables explicit derivation of the null distribution of the test statistic, allowing fast and statistically rigorous significance testing. Consequently, CONCISE achieves both computational efficiency and robust statistical performance.

Beyond its methodological and performance advances, the applications of CONCISE have also generated biologically meaningful insights across diverse biological systems, including developing mouse embryo, mouse model with inflammatory bowel diseases, human breast cancer and non-small cell lung cancer tissues. Particularly, it has characterized disease-associated CCC. For instance, it revealed multiple unique communication patterns originating from inflammation-associated fibroblasts that are distinct from normal fibroblasts in intestine inflammation. It also delineated extensive tumor-immune and tumor-stromal signaling, corresponding to immune suppression, stromal activation, angiogenesis and tumor progression in tumor microenvironment.

One potential limitation of the current CONCISE framework, despite its advantages, is that it primarily focuses on the inference of spatial LRIs and does not incorporate downstream transcriptional responses induced by such interactions. Although L-R co-expression provides direct evidence of potential cellular communication, integrating information from downstream target genes and signaling programs may further strengthen the evidence for functional interactions, improve statistical power, and enhance biological interpretability. Extending CONCISE to incorporate downstream signaling programs and gene regulatory responses represents an important direction for future methodological development.

Reliable inference of CCC is essential for understanding how multicellular systems are organized, maintained, and remodeled during development, homeostasis, and disease. As increasingly diverse spatial transcriptomics datasets capturing rich biological contexts continue to accumulate, the need for statistically rigorous and efficient approaches to CCC inference will only grow. We expect that CONCISE, with its exceptional performance, will serve as a valuable addition to the spatial transcriptomics analytical toolkit and facilitate more reliable characterization of cellular communication across a broad range of biological systems and disease contexts.

## Methods

### The model of CONCISE

To infer cell-cell communication from spatial transcriptomics data, CONCISE aims to identify ligand-receptor pairs showing significant spatial co-expression from a comprehensive candidate set. By default, it uses ligand-receptor lists from CellChatDB [14] as input, while users may adopt customized lists.

To ensure reliable detection of spatial co-expression between ligand-receptor pairs, we propose an expression-measurement model that simultaneously accounts for multiple confounding factors, including varying total molecular counts across spots or cells, measurement errors, and spatial autocorrelation in spatially resolved gene expression, all of which, when ignored, may lead to spurious results.

Let *i* = 1, 2, · · ·, *n* index spatial spots or cells in the analyzed spatial transcriptomics section. The spatial location of spot or cell *i* is denoted by **t**_*i*_ = (*t*_*i*1_, *t*_*i*2_) ∈ ℝ^2^, and we collect the spatial locations of all spots or cells in **t** = (**t**_1_, · · ·, **t**_*n*_)^*T*^. For spot or cell *i*, its observed gene expression measurements are represented by (*y*_1*i*_, · · ·, *y*_*gi*_) ∈ ℕ^*g*^, where we use *y*_*pi*_ to denote the observed count for gene *p* in spot or cell *i*. The total molecular counts of spot or cell *i* is defined as

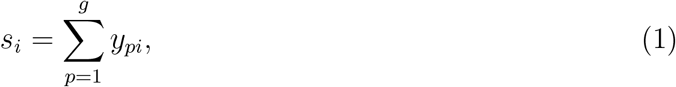

which is the sum of observed counts across all measured genes *p* = 1, · · ·, *g* in this spot or cell. For inferring ligand-receptor spatial co-expression, we denote the observed counts for ligand *p*_1_ and receptor *p*_2_ across all *n* spots or cells by 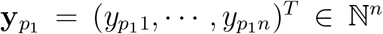 and 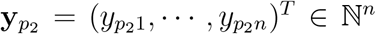, respectively. Similarly, let 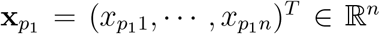 and 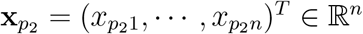 represent the corresponding underlying expression levels, defined as the number of molecules from ligand *p*_1_ and target *p*_2_ relative to the total number of molecules in each spot or cell. We model the observed counts 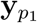 and 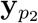 as:

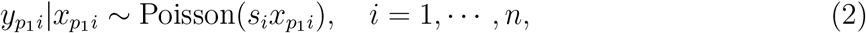

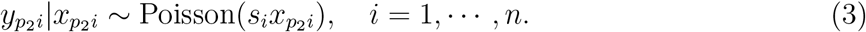

Here, the measured counts 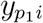 and 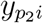 of ligand *p*_1_ and receptor *p*_2_ in spot or cell *i* are assumed to follow a Poisson measurement model [34, 40] depending on the underlying expression levels 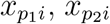, and total molecular counts *s*_*i*_. This Poisson measurement model explicitly accounts for the total molecular counts and measurement errors [40, 115, 116, 117, 118, 119].

In the context of spatial transcriptomics data, the unknown underlying expression levels 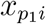 and 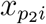 can be viewed as the unobserved spatial random process occurring at location **t**_*i*_, corresponding to spot or cell *i* [40, 120]. To infer the underlying spatial co-expression between ligand *p*_1_ and receptor *p*_2_, we model latent variables 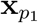 and 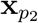 using the following spatial process framework:

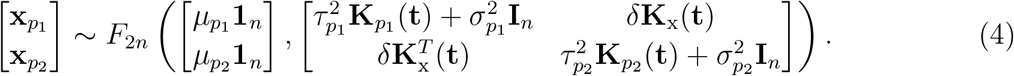

where *F*_2*n*_(***µ*, Σ**) denotes an unknown nonnegative 2*n*-variate distribution with mean vector ***µ*** and covariance matrix **Σ**, vector **1** = (1, · · ·, 1)^*T*^ ∈ ℝ^*n*^, and **I**_*n*_ is an *n*-dimensional identify matrix. In this model, 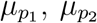 are the mean expression levels of ligand *p*_1_ and receptor *p*_2_, respectively. 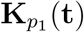 and 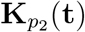 are spatial kernel functions, with the (*i, j*)th entry given by 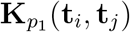 and 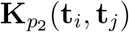 We normalize matrices 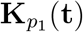 and 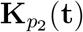, so that their diagonal elements equal one. According to Eq. (4), marginal models for underlying expression levels 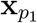 and 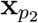 are given by the following spatial processes:

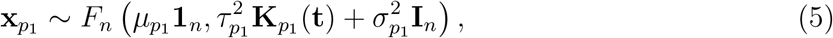

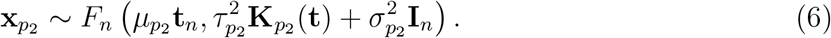

In spatial statistics, scaling factors 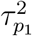 and 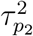 are commonly known to quantify the expression variance in 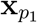 and 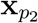 captured by spatial patterns or spatial location information, whereas scaling factors 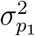 and 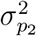 measure the variance arising from random noise independent of spatial locations. The spatial process model in Eqs. (4)-(6), incorporating spatial kernels 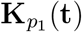 and 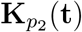, explicitly account for spatial autocorrelation in the data, which can bias the estimation of ligand-receptor co-expression and lead to false positives if not properly addressed [26, 27]. With spatial autocorrelation in both 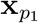 and 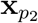 accounted for, we estimate their spatial co-expression through parameter *δ* in Eq. (4).

CONCISE accommodates two complementary forms of spatial co-expression analysis. The first evaluates co-expression between spatially neighboring spots or cells and forms the basis of spatial LRI inference. The second evaluates co-expression within the same spot or cell and can be used to construct spatial gene-gene co-expression networks. These two settings are unified within a common statistical framework through the specification of the interaction kernel **K**_x_(**t**) in Eq. (4). For spatial LRI inference, **K**_x_(**t**) is defined according to spatial proximity, with **K**_x_(**t**_*i*_, **t**_*j*_) = 1 if locations *i* and *j* are spatial neighbors and **K**_x_(**t**_*i*_, **t**_*j*_) = 0 otherwise. In contrast, setting **K**_x_(**t**) = **I**_*n*_ restricts co-expression assessment to measurements obtained from the same spatial location, thereby recovering within-location co-expression analysis for spatial gene-gene network construction.

### Parameter estimation in CONCISE

To infer spatial co-expression between ligand *p*_1_ and receptor *p*_2_ based on the statistical model with Eqs. (2)-(4), we estimate unknown parameters 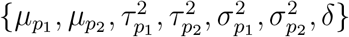. Under the Poisson measurement model, for *q* ∈ *{p*_1_, *p*_2_*}* it holds that 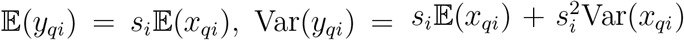, and for *q*_1_, *q*_2_ ∈ *{p*_1_, *p*_2_*}* with *q*_1_ ≠ *q*_2_ or *i* ≠ *j*, 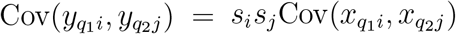. These relationships motivate a moment-based approach for parameter estimation, without imposing distributional assumptions on *F*_2*n*_ [35].

For notation convenience, we collect the total molecular counts of all spots or cells in **s** = (*s*, · · ·, *s*)^*T*^ ∈ ℕ^*n*^. The mean expression levels 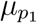 and 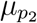 can be estimated by 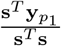 and 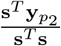, respectively. For variance and covariance estimation, we denote 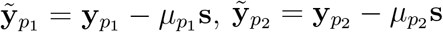, and 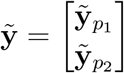. The second-order moment of 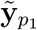 and 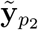 for ligand *p*_1_ and receptor *p*_2_ is then given by:

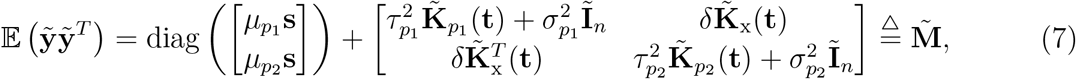

where

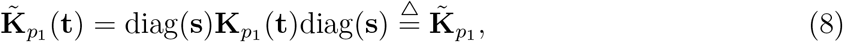

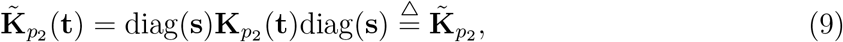

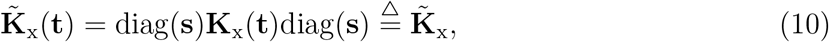

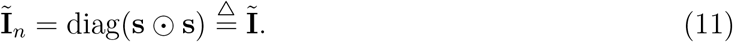

We estimate the parameters by matching the empirical and theoretical second-order moments using the least-squares criterion [121, 122]: 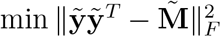. Full details of the derivation are provided in Supplementary Note 1, leading to the following estimating equations:

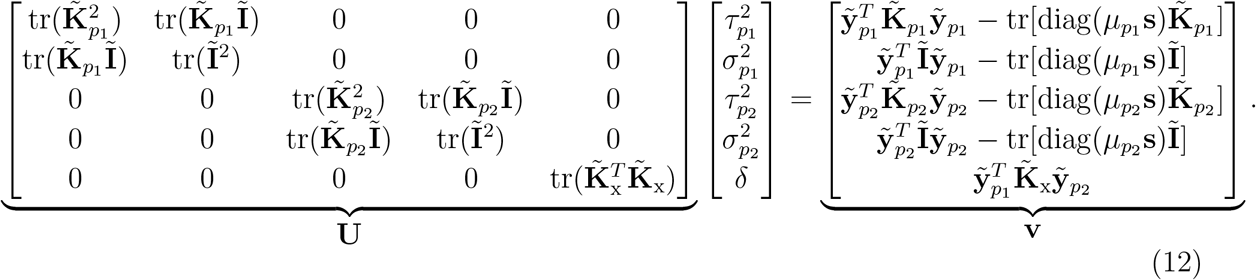

In Eq. (12), the matrix **U** needs to be computed only once for the analyzed spatial transcriptomics section, while for each ligand-receptor pair, only the vector **v** must be recalculated.

### Statistical inference in CONCISE

Next, we develop a statistical test to assess whether the expression levels of a ligand–receptor pair are independent in the analyzed spatial transcriptomics section. Independence would imply no interaction between them, corresponding to the null hypothesis *H*_0_ : *δ* = 0, while the alternative is *H*_1_ : *δ* ≠ 0.

From Eq. (12), the core parameter *δ*, which quantifies the spatial co-expression between a ligand–receptor pair, is estimated by

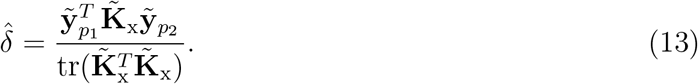

The variance of the numerator, 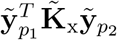, can be derived under the statistical model specified in Eqs. (2)–(4). The resulting expression is

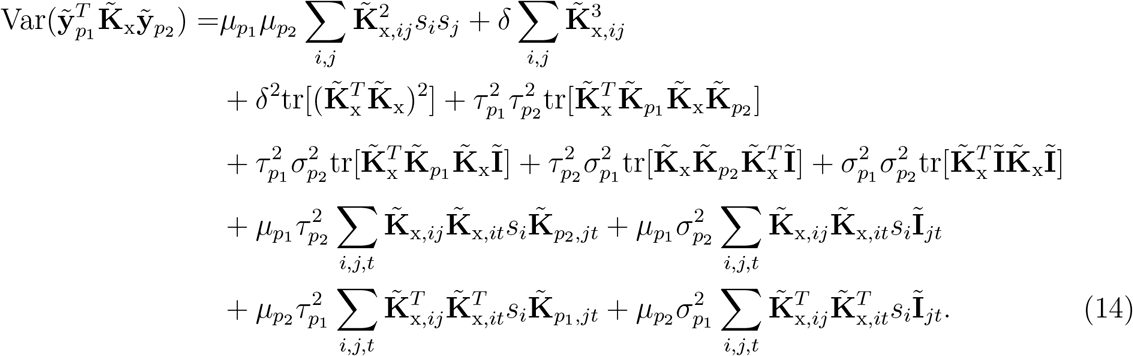

Full details of this derivation are provided in Supplementary Note 2.

To perform hypothesis testing, we define the test statistic

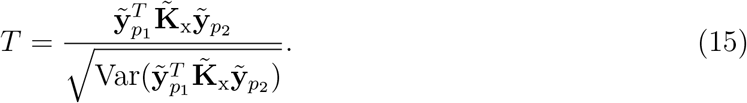

Under the null hypothesis *H*_0_ : *δ* = 0, the statistic *T* follows *T* ~ *N* (0, 1). This enables direct computation of the *p*-value with the estimated parameters 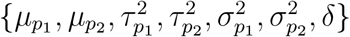.

In the derived expression of 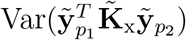, Eq. (14), the terms 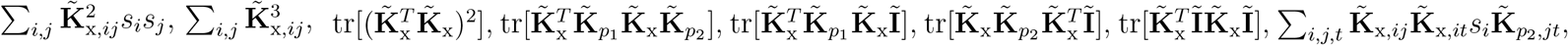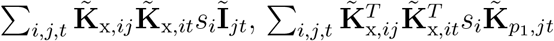, and 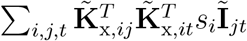 depend only on the total molecular counts and spatial coordinates of spots or cells in the analyzed tissue section. These quantities therefore need to be computed only once, which reduces the computational cost of testing multiple ligand-receptor pairs.

## Data availability

All data used in this work are publicly available through online sources.

- Human breast cancer dataset profiled by Visium platform [36] (https://www.10xgenomics.com/datasets/human-breast-cancer-block-a-section-1-1-standard-1-0-0).
- Mouse embryo dataset profiled by Stereo-seq [37] (https://db.cngb.org/stomics/mosta/).
- Mouse intestinal inflammation dataset profiled by MERFISH [38] (https://doi.org/10.5061/dryad.rjdfn2zh3).
- Human NSCLC dataset profiled by CosMx [86] (https://brukerspatialbiology.com/products/cosmx-spatial-molecular-imager/ffpe-dataset/).

## Code availability

The CONCISE software is available at https://github.com/jiazhao97/CONCISE.

## Acknowledgements

This work was supported in part by NIH grants P50CA196530, U24HG012108, U01HG013840, and R01DA065184 to H.Z.

## Notes

### Competing Interest Statement

The authors have declared no competing interest.

