## Supplementary Information for "Spatial co-expression and cell-cell communication inference from spatially resolved transcriptomics with CONCISE"

### Supplementary Notes

#### Supplementary Note 1: Parameter estimation

Note that the second-order moment of $\tilde{\mathbf{y}}_{p_1}$ and $\tilde{\mathbf{y}}_{p_2}$ for ligand $p_1$ and receptor $p_2$ is given by Eq. (7). The moment-based estimator is therefore obtained by matching the empirical and theoretical second-order moments under the least-squares criterion:

$$\text{argmin}_{\tau_{p_1}^2,\tau_{p_2}^2,\sigma_{p_1}^2,\sigma_{p_2}^2,\delta}\, \|\tilde{\mathbf{y}}\tilde{\mathbf{y}}^T - \tilde{\mathbf{M}}\|_F^2. \tag{1}$$

For notation convenience, let

$$\tilde{\boldsymbol{\mu}}_{p_1} = \mu_{p_1}\mathbf{s}, \quad \tilde{\boldsymbol{\mu}}_{p_2} = \mu_{p_2}\mathbf{s}, \quad \tilde{\boldsymbol{\mu}} = \begin{bmatrix}\tilde{\boldsymbol{\mu}}_{p_1}\\ \tilde{\boldsymbol{\mu}}_{p_2}\end{bmatrix}, \tag{2}$$

Then,

$$\tilde{\mathbf{M}} = \text{diag}(\tilde{\boldsymbol{\mu}}) + \begin{bmatrix}\tau_{p_1}^2\tilde{\mathbf{K}}_{p_1} & \delta\tilde{\mathbf{K}}_{\text{x}}\\ \delta\tilde{\mathbf{K}}_{\text{x}}^T & \tau_{p_2}^2\tilde{\mathbf{K}}_{p_2}\end{bmatrix} + \begin{bmatrix}\sigma_{p_1}^2\tilde{\mathbf{I}} & \mathbf{0}\\ \mathbf{0} & \sigma_{p_2}^2\tilde{\mathbf{I}}\end{bmatrix}. \tag{3}$$

Using $\|\mathbf{B}\|_F = \sqrt{\mathrm{tr}(\mathbf{B}\mathbf{B}^T)}$, the objective function in Supplementary Eq. (1) expands to:

$$
\begin{aligned}
&\|\tilde{\mathbf{y}}\tilde{\mathbf{y}}^T - \tilde{\mathbf{M}}\|_F^2 \\
=&\mathrm{tr}\left\{\begin{bmatrix}\tau_{p_1}^2\tilde{\mathbf{K}}_{p_1} & \delta\tilde{\mathbf{K}}_{\mathrm{x}} \\ \delta\tilde{\mathbf{K}}_{\mathrm{x}}^T & \tau_{p_2}^2\tilde{\mathbf{K}}_{p_2}\end{bmatrix}\begin{bmatrix}\tau_{p_1}^2\tilde{\mathbf{K}}_{p_1} & \delta\tilde{\mathbf{K}}_{\mathrm{x}} \\ \delta\tilde{\mathbf{K}}_{\mathrm{x}}^T & \tau_{p_2}^2\tilde{\mathbf{K}}_{p_2}\end{bmatrix}^T\right\} + \mathrm{tr}\left\{\begin{bmatrix}\sigma_{p_1}^2\tilde{\mathbf{I}} & \mathbf{0} \\ \mathbf{0} & \sigma_{p_2}^2\tilde{\mathbf{I}}\end{bmatrix}\begin{bmatrix}\sigma_{p_1}^2\tilde{\mathbf{I}} & \mathbf{0} \\ \mathbf{0} & \sigma_{p_2}^2\tilde{\mathbf{I}}\end{bmatrix}^T\right\} \\
&- 2\mathrm{tr}\left\{\begin{bmatrix}\tilde{\mathbf{y}}_{p_1}\tilde{\mathbf{y}}_{p_1}^T - \mathrm{diag}(\tilde{\boldsymbol{\mu}}_{p_1}) & \tilde{\mathbf{y}}_{p_1}\tilde{\mathbf{y}}_{p_2}^T \\ \tilde{\mathbf{y}}_{p_2}\tilde{\mathbf{y}}_{p_1}^T & \tilde{\mathbf{y}}_{p_2}\tilde{\mathbf{y}}_{p_2}^T - \mathrm{diag}(\tilde{\boldsymbol{\mu}}_{p_2})\end{bmatrix}\begin{bmatrix}\tau_{p_1}^2\tilde{\mathbf{K}}_{p_1} & \delta\tilde{\mathbf{K}}_{\mathrm{x}} \\ \delta\tilde{\mathbf{K}}_{\mathrm{x}}^T & \tau_{p_2}^2\tilde{\mathbf{K}}_{p_2}\end{bmatrix}^T\right\} \\
&- 2\mathrm{tr}\left\{\begin{bmatrix}\tilde{\mathbf{y}}_{p_1}\tilde{\mathbf{y}}_{p_1}^T - \mathrm{diag}(\tilde{\boldsymbol{\mu}}_{p_1}) & \tilde{\mathbf{y}}_{p_1}\tilde{\mathbf{y}}_{p_2}^T \\ \tilde{\mathbf{y}}_{p_2}\tilde{\mathbf{y}}_{p_1}^T & \tilde{\mathbf{y}}_{p_2}\tilde{\mathbf{y}}_{p_2}^T - \mathrm{diag}(\tilde{\boldsymbol{\mu}}_{p_2})\end{bmatrix}\begin{bmatrix}\sigma_{p_1}^2\tilde{\mathbf{I}} & \mathbf{0} \\ \mathbf{0} & \sigma_{p_2}^2\tilde{\mathbf{I}}\end{bmatrix}^T\right\} \\
&+ 2\mathrm{tr}\left\{\begin{bmatrix}\tau_{p_1}^2\tilde{\mathbf{K}}_{p_1} & \delta\tilde{\mathbf{K}}_{\mathrm{x}} \\ \delta\tilde{\mathbf{K}}_{\mathrm{x}}^T & \tau_{p_2}^2\tilde{\mathbf{K}}_{p_2}\end{bmatrix}\begin{bmatrix}\sigma_{p_1}^2\tilde{\mathbf{I}} & \mathbf{0} \\ \mathbf{0} & \sigma_{p_2}^2\tilde{\mathbf{I}}\end{bmatrix}^T\right\} \\
&+ \mathrm{tr}\left\{(\tilde{\mathbf{y}}\tilde{\mathbf{y}}^T - \mathrm{diag}(\tilde{\boldsymbol{\mu}}))(\tilde{\mathbf{y}}\tilde{\mathbf{y}}^T - \mathrm{diag}(\tilde{\boldsymbol{\mu}}))^T\right\}
\end{aligned}
$$

$$
\begin{aligned}
=&(\tau_{p_1}^2)^2\mathrm{tr}\left(\tilde{\mathbf{K}}_{p_1}^2\right) + (\tau_{p_2}^2)^2\mathrm{tr}\left(\tilde{\mathbf{K}}_{p_2}^2\right) + 2\delta^2\mathrm{tr}\left(\tilde{\mathbf{K}}_{\mathrm{x}}^T\tilde{\mathbf{K}}_{\mathrm{x}}\right) + (\sigma_{p_1}^2)^2\mathrm{tr}\left(\tilde{\mathbf{I}}^2\right) + (\sigma_{p_2}^2)^2\mathrm{tr}\left(\tilde{\mathbf{I}}^2\right) \\
&- 2\tau_{p_1}^2\left\{\tilde{\mathbf{y}}_{p_1}^T\tilde{\mathbf{K}}_{p_1}\tilde{\mathbf{y}}_{p_1} - \mathrm{tr}\left[\mathrm{diag}(\tilde{\boldsymbol{\mu}}_{p_1})\tilde{\mathbf{K}}_{p_1}\right]\right\} - 2\tau_{p_2}^2\left\{\tilde{\mathbf{y}}_{p_2}^T\tilde{\mathbf{K}}_{p_2}\tilde{\mathbf{y}}_{p_2} - \mathrm{tr}\left[\mathrm{diag}(\tilde{\boldsymbol{\mu}}_{p_2})\tilde{\mathbf{K}}_{p_2}\right]\right\} \\
&- 4\delta\tilde{\mathbf{y}}_{p_1}^T\tilde{\mathbf{K}}_{\mathrm{x}}\tilde{\mathbf{y}}_{p_2} - 2\sigma_{p_1}^2\left\{\tilde{\mathbf{y}}_{p_1}^T\tilde{\mathbf{I}}\tilde{\mathbf{y}}_{p_1} - \mathrm{tr}\left[\mathrm{diag}(\tilde{\boldsymbol{\mu}}_{p_1})\tilde{\mathbf{I}}\right]\right\} - 2\sigma_{p_2}^2\left\{\tilde{\mathbf{y}}_{p_2}^T\tilde{\mathbf{I}}\tilde{\mathbf{y}}_{p_2} - \mathrm{tr}\left[\mathrm{diag}(\tilde{\boldsymbol{\mu}}_{p_2})\tilde{\mathbf{I}}\right]\right\} \\
&+ 2\tau_{p_1}^2\sigma_{p_1}^2\mathrm{tr}\left(\tilde{\mathbf{K}}_{p_1}\tilde{\mathbf{I}}\right) + 2\tau_{p_2}^2\sigma_{p_2}^2\mathrm{tr}\left(\tilde{\mathbf{K}}_{p_2}\tilde{\mathbf{I}}\right) + \mathrm{tr}\left\{(\tilde{\mathbf{y}}\tilde{\mathbf{y}}^T - \mathrm{diag}(\tilde{\boldsymbol{\mu}}))(\tilde{\mathbf{y}}\tilde{\mathbf{y}}^T - \mathrm{diag}(\tilde{\boldsymbol{\mu}}))^T\right\}. \qquad (4)
\end{aligned}
$$

Taking derivatives of the objective function with respect to $\{\tau_{p_1}^2, \tau_{p_2}^2, \sigma_{p_1}^2, \sigma_{p_2}^2, \delta\}$ and setting them to zero leads to the estimating equation given by Eq. (12).

### Supplementary Note 2: Statistical inference

According to Eq. (12) derived for parameter estimation, the core parameter $\delta$, which quantifies the spatial co-expression between a ligand–receptor pair, is estimated as:

$$\hat{\delta} = \frac{\tilde{\mathbf{y}}_{p_1}^T\tilde{\mathbf{K}}_{\mathrm{x}}\tilde{\mathbf{y}}_{p_2}}{\mathrm{tr}(\tilde{\mathbf{K}}_{\mathrm{x}}^T\tilde{\mathbf{K}}_{\mathrm{x}})}. \qquad (5)$$

To perform hypothesis testing, we need to derive the variance of this estimator under the statistical model specified in Eqs. (2)-(4).

Let the matrix $\mathbf{A}$ be defined as

$$\mathbf{A} = \begin{bmatrix}\mathbf{0} & \tilde{\mathbf{K}}_{\mathrm{x}} \\ \tilde{\mathbf{K}}_{\mathrm{x}}^T & \mathbf{0}\end{bmatrix}. \qquad (6)$$

Then, the variance of $\tilde{\mathbf{y}}_{p_1}^T\tilde{\mathbf{K}}_2\tilde{\mathbf{y}}_{p_2}$ can be expressed as

$$\mathrm{Var}(\tilde{\mathbf{y}}_{p_1}^T\tilde{\mathbf{K}}_{\mathrm{x}}\tilde{\mathbf{y}}_{p_2}) = \frac{1}{4}\mathrm{Var}\{\tilde{\mathbf{y}}^T\mathbf{A}\tilde{\mathbf{y}}\} = \frac{1}{4}\left\{\mathbb{E}\left[(\tilde{\mathbf{y}}^T\mathbf{A}\tilde{\mathbf{y}})^2\right] - \left[\mathbb{E}(\tilde{\mathbf{y}}^T\mathbf{A}\tilde{\mathbf{y}})\right]^2\right\}. \qquad (7)$$

In the following, we derive $\mathbb{E}(\tilde{\mathbf{y}}^T\mathbf{A}\tilde{\mathbf{y}})$ and $\mathbb{E}[(\tilde{\mathbf{y}}^T\mathbf{A}\tilde{\mathbf{y}})^2]$, respectively.

**Derivation of $\mathbb{E}(\tilde{\mathbf{y}}^T\mathbf{A}\tilde{\mathbf{y}})$.**

For simplicity, we define

$$\mathbf{S} = \text{diag}\left(\begin{bmatrix}\mathbf{s}\\ \mathbf{s}\end{bmatrix}\right), \quad \boldsymbol{\Sigma} = \begin{bmatrix}\tau_{p_1}^2\mathbf{K}_{p_1}(\mathbf{t}) + \sigma_{p_1}^2\mathbf{I}_n & \delta\mathbf{K}_{\text{x}}(\mathbf{t})\\ \delta\mathbf{K}_{\text{x}}^T(\mathbf{t}) & \tau_{p_2}^2\mathbf{K}_{p_2}(\mathbf{t}) + \sigma_{p_2}^2\mathbf{I}_n\end{bmatrix}, \quad \boldsymbol{\mu} = \begin{bmatrix}\mu_{p_1}\mathbf{1}\\ \mu_{p_2}\mathbf{1}\end{bmatrix}. \tag{8}$$

From the statistical model in Eqs. (2)-(4), we have

$$\text{Var}(\tilde{\mathbf{y}}) = \mathbf{S}\text{diag}(\boldsymbol{\mu}) + \mathbf{S}\boldsymbol{\Sigma}\mathbf{S}. \tag{9}$$

Therefore,

$$\mathbb{E}(\tilde{\mathbf{y}}^T\mathbf{A}\tilde{\mathbf{y}}) = \mathbb{E}[\text{tr}(\mathbf{A}\tilde{\mathbf{y}}\tilde{\mathbf{y}}^T)] = \text{tr}[\mathbf{A}\text{Var}(\tilde{\mathbf{y}})] = \text{tr}(\mathbf{A}\mathbf{S}\boldsymbol{\Sigma}\mathbf{S}), \tag{10}$$

since $\text{tr}[\mathbf{A}\mathbf{S}\text{diag}(\boldsymbol{\mu})] = 0$.

**Derivation of $\mathbb{E}[(\tilde{\mathbf{y}}^T\mathbf{A}\tilde{\mathbf{y}})^2]$.**

For simplicity, we define

$$\mathbf{y} = \begin{bmatrix}\mathbf{y}_{p_1}\\ \mathbf{y}_{p_2}\end{bmatrix}, \quad \mathbf{x} = \begin{bmatrix}\mathbf{x}_{p_1}\\ \mathbf{x}_{p_2}\end{bmatrix}, \quad \boldsymbol{\epsilon} = \mathbf{y} - \mathbf{S}\mathbf{x}, \quad \mathbf{z} = \mathbf{S}(\mathbf{x} - \boldsymbol{\mu}), \tag{11}$$

so that

$$\tilde{\mathbf{y}} = \boldsymbol{\epsilon} + \mathbf{z}. \tag{12}$$

Here, $\boldsymbol{\epsilon}$ represents the Poisson measurement noise conditional on $\mathbf{x}$, as described in Eqs. (2) and (3), while the distributional properties of $\mathbf{z}$ follow from the spatial process model in Eq. (4). Expanding $\mathbb{E}[(\tilde{\mathbf{y}}^T\mathbf{A}\tilde{\mathbf{y}})^2]$ using Supplementary Eq. (12) gives:

$$\begin{aligned}\mathbb{E}[(\tilde{\mathbf{y}}^T\mathbf{A}\tilde{\mathbf{y}})^2] =& \mathbb{E}[(\boldsymbol{\epsilon}^T\mathbf{A}\boldsymbol{\epsilon})^2] + \mathbb{E}[(\mathbf{z}^T\mathbf{A}\mathbf{z})^2] + 4\mathbb{E}[(\boldsymbol{\epsilon}^T\mathbf{A}\mathbf{z})^2]\\ &+ 2\mathbb{E}[\boldsymbol{\epsilon}^T\mathbf{A}\boldsymbol{\epsilon}\mathbf{z}^T\mathbf{A}\mathbf{z}] + 4\mathbb{E}[\boldsymbol{\epsilon}^T\mathbf{A}\boldsymbol{\epsilon}\boldsymbol{\epsilon}^T\mathbf{A}\mathbf{z}] + 4\mathbb{E}[\mathbf{z}^T\mathbf{A}\mathbf{z}\boldsymbol{\epsilon}^T\mathbf{A}\mathbf{z}].\end{aligned} \tag{13}$$

We next derive each of these terms under the null hypothesis.

<u>Derivation of $\mathbb{E}[(\boldsymbol{\epsilon}^T\mathbf{A}\boldsymbol{\epsilon})^2]$</u>.

For notational convenience, we introduce a global index $k = 1, \cdots, 2n$ to refer to elements of the $2n$-dimensional vectors $\mathbf{y}$, $\mathbf{s}$, $\mathbf{x}$, $\boldsymbol{\epsilon}$, $\mathbf{z}$, $\boldsymbol{\mu}$ and $\tilde{\boldsymbol{\mu}}$. Likewise, we use global indices $k, l = 1, \cdots, 2n$ to denote elements of the $2n \times 2n$ matrices $\boldsymbol{\Sigma}$ and $\mathbf{A}$.

Under the Poisson measurement model in Eqs. (2) and (3), for any $k, l \in \{1, \cdots, 2n\}$ with $k \neq l$, the noise terms $\epsilon_k$ and $\epsilon_l$ are conditionally independent given $\mathbf{x}$. Moreover,

$$\mathbb{E}(\epsilon_k|\mathbf{x}) = 0, \quad \mathbb{E}(\epsilon_k^2|\mathbf{x}) = s_k x_k, \quad k = 1, \cdots, 2n. \tag{14}$$

From the spatial process model in Eq. (4), for $k, l \in \{1, \cdots, 2n\}$ with $k \neq l$, we have

$$\mathbb{E}(x_k x_l) = \mu_k \mu_l + \Sigma_{kl}. \tag{15}$$

Since $\mathbf{A}$ is symmetric and $\mathbf{A}_{kk} = 0$, it follows that

$$\boldsymbol{\epsilon}^T \mathbf{A} \boldsymbol{\epsilon} = 2 \sum_{k<l} A_{kl} \epsilon_k \epsilon_l. \tag{16}$$

Therefore,

$$\begin{aligned}
\mathbb{E}[(\boldsymbol{\epsilon}^T \mathbf{A} \boldsymbol{\epsilon})^2] =& \mathbb{E}_{\mathbf{x}} \mathbb{E} \left[ 4 \sum_{k<l} \sum_{u<v} A_{kl} A_{uv} \epsilon_k \epsilon_l \epsilon_u \epsilon_v | \mathbf{x} \right] \\
=& \mathbb{E}_{\mathbf{x}} \mathbb{E} \left[ 4 \sum_{k<l} A_{kl}^2 \epsilon_k^2 \epsilon_l^2 | \mathbf{x} \right] \\
=& \mathbb{E} \left[ 4 \sum_{k<l} A_{kl}^2 s_k x_k s_l x_l \right] \\
=& 4 \sum_{k<l} A_{kl}^2 s_k s_l \left( \mu_k \mu_l + \Sigma_{kl} \right).
\end{aligned} \tag{17}$$

<u>Derivation of $\mathbb{E}[(\mathbf{z}^T \mathbf{A} \mathbf{z})^2]$</u>.

From the spatial process model in Eq. 4, we have

$$\text{Var}(\mathbf{z}) = \mathbf{S} \boldsymbol{\Sigma} \mathbf{S}, \tag{18}$$

$$\mathbb{E}(\mathbf{z}^T \mathbf{A} \mathbf{z}) = \mathbb{E}[\text{tr}(\mathbf{A} \mathbf{z} \mathbf{z}^T)] = \text{tr}[\mathbf{A} \text{Var}(\mathbf{z})] = \text{tr}(\mathbf{A} \mathbf{S} \boldsymbol{\Sigma} \mathbf{S}). \tag{19}$$

Approximating $\text{Var}(\mathbf{z}^T \mathbf{A} \mathbf{z})$ by

$$\text{Var}(\mathbf{z}^T \mathbf{A} \mathbf{z}) = 2\text{tr}[(\mathbf{A} \mathbf{S} \boldsymbol{\Sigma} \mathbf{S})^2], \tag{20}$$

we obtain

$$\begin{aligned}
\mathbb{E}[(\mathbf{z}^T \mathbf{A} \mathbf{z})^2] &= \text{Var}(\mathbf{z}^T \mathbf{A} \mathbf{z}) + [\mathbb{E}(\mathbf{z}^T \mathbf{A} \mathbf{z})]^2 \\
&= 2\text{tr}[(\mathbf{A} \mathbf{S} \boldsymbol{\Sigma} \mathbf{S})^2] + [\text{tr}(\mathbf{A} \mathbf{S} \boldsymbol{\Sigma} \mathbf{S})]^2.
\end{aligned} \tag{21}$$

<u>Derivation of $\mathbb{E}[(\boldsymbol{\epsilon}^T \mathbf{A} \mathbf{z})^2]$</u>.

With $\mathbf{A}_{kk} = 0$, the term $\boldsymbol{\epsilon}^T \mathbf{A} \mathbf{z}$ can be written as

$$\boldsymbol{\epsilon}^T \mathbf{A} \mathbf{z} = \sum_{k<l} A_{kl} \epsilon_k z_l + \sum_{k>l} A_{kl} \epsilon_k z_l. \tag{22}$$

Therefore,

$$(\boldsymbol{\epsilon}^T\mathbf{A}\mathbf{z})^2 = \left(\sum_{k<l} A_{kl}\epsilon_k z_l\right)^2 + \left(\sum_{k>l} A_{kl}\epsilon_k z_l\right)^2 + 2\left(\sum_{k<l} A_{kl}\epsilon_k z_l\right)\left(\sum_{k>l} A_{kl}\epsilon_k z_l\right). \tag{23}$$

From the Poisson measurement model in Eqs. (2) and (3), for $k \neq l$, the noise terms $\epsilon_k$ and $\epsilon_l$ are conditionally independent given $\mathbf{x}$, and satisfy $\mathbb{E}(\epsilon_k|\mathbf{x}) = 0$, $\mathbb{E}(\epsilon_k^2|\mathbf{x}) = s_k x_k$. Using these properties, we have

$$\begin{aligned}
\mathbb{E}[(\boldsymbol{\epsilon}^T\mathbf{A}\mathbf{z})^2] =& \mathbb{E}_{\mathbf{x}}\mathbb{E}\left[\sum_{k<l} A_{kl}^2\epsilon_k^2 z_l^2 + 2\sum_{k<l<u} A_{kl}A_{ku}\epsilon_k^2 z_l z_u|\mathbf{x}\right] \\
&+ \mathbb{E}_{\mathbf{x}}\mathbb{E}\left[\sum_{k>l} A_{kl}^2\epsilon_k^2 z_l^2 + 2\sum_{k>l>u} A_{kl}A_{ku}\epsilon_k^2 z_l z_u|\mathbf{x}\right] + \mathbb{E}_{\mathbf{x}}\mathbb{E}\left[2\sum_{l<k<u} A_{kl}A_{ku}\epsilon_k^2 z_l z_u|\mathbf{x}\right] \\
=& \mathbb{E}\left[\sum_{k<l} A_{kl}^2 s_k x_k s_l^2(x_l-\mu_l)^2 + 2\sum_{k<l<u} A_{kl}A_{ku}s_k x_k s_l(x_l-\mu_l)s_u(x_u-\mu_u)\right] \\
&+ \mathbb{E}\left[\sum_{k>l} A_{kl}^2 s_k x_k s_l^2(x_l-\mu_l)^2 + 2\sum_{k>l>u} A_{kl}A_{ku}s_k x_k s_l(x_l-\mu_l)s_u(x_u-\mu_u)\right] \\
&+ \mathbb{E}\left[2\sum_{l<k<u} A_{kl}A_{ku}s_k x_k s_l(x_l-\mu_l)s_u(x_u-\mu_u)\right].
\end{aligned} \tag{24}$$

Assuming that the third central moment of $\mathbf{x}$ can be neglected in computing the variance of $\tilde{\mathbf{y}}_{p_1}^T\tilde{\mathbf{K}}_{\mathrm{x}}\tilde{\mathbf{y}}_{p_2}$, and noting from Supplementary Eq. (6) that $A_{kl}A_{ku} = 0$ whenever $l < k < u$, we obtain $\mathbb{E}[(\boldsymbol{\epsilon}^T\mathbf{A}\mathbf{z})^2]$ as

$$\begin{aligned}
\mathbb{E}[(\boldsymbol{\epsilon}^T\mathbf{A}\mathbf{z})^2] =& \sum_{k<l} A_{kl}^2 s_k s_l^2\mu_k\Sigma_{ll} + 2\sum_{k<l<u} A_{kl}A_{ku}s_k s_l s_u\mu_k\Sigma_{lu} \\
&+ \sum_{k>l} A_{kl}^2 s_k s_l^2\mu_k\Sigma_{ll} + 2\sum_{k>l>u} A_{kl}A_{ku}s_k s_l s_u\mu_k\Sigma_{lu} \\
=& \sum_{k<l,k<u} A_{kl}A_{ku}s_k\mu_k s_l s_u\Sigma_{lu} + \sum_{k>l,k>u} A_{kl}A_{ku}s_k\mu_k s_l s_u\Sigma_{lu}.
\end{aligned} \tag{25}$$

<u>Derivation of $\mathbb{E}[\boldsymbol{\epsilon}^T\mathbf{A}\boldsymbol{\epsilon}\mathbf{z}^T\mathbf{A}\mathbf{z}]$, $\mathbb{E}[\boldsymbol{\epsilon}^T\mathbf{A}\boldsymbol{\epsilon}\boldsymbol{\epsilon}^T\mathbf{A}\mathbf{z}]$ and $\mathbb{E}[\mathbf{z}^T\mathbf{A}\mathbf{z}\boldsymbol{\epsilon}^T\mathbf{A}\mathbf{z}]$</u>.

Since for any $k, l = 1, \cdots, 2n$ with $k \neq l$, the noise terms $\epsilon_k$ and $\epsilon_l$ are conditionally independent given $\mathbf{x}$, and $\mathbb{E}(\epsilon_k|\mathbf{x}) = 0$, it follows that

$$\mathbb{E}[\boldsymbol{\epsilon}^T\mathbf{A}\boldsymbol{\epsilon}\mathbf{z}^T\mathbf{A}\mathbf{z}] = \mathbb{E}_{\mathbf{x}}\mathbb{E}\left[4\left(\sum_{k<l} A_{kl}\epsilon_k\epsilon_l\right)\left(\sum_{k<l} A_{kl}z_k z_l\right)|\mathbf{x}\right] = 0, \tag{26}$$

$$\mathbb{E}[\boldsymbol{\epsilon}^T\mathbf{A}\boldsymbol{\epsilon}\boldsymbol{\epsilon}^T\mathbf{A}\mathbf{z}] = \mathbb{E}_{\mathbf{x}}\mathbb{E}\left[2\left(\sum_{k<l} A_{kl}\epsilon_k\epsilon_l\right)\left(\sum_{k,l} A_{kl}\epsilon_k z_l\right)|\mathbf{x}\right] = 0, \tag{27}$$

$$\mathbb{E}[\mathbf{z}^T\mathbf{A}\mathbf{z}\boldsymbol{\epsilon}^T\mathbf{A}\mathbf{z}] = \mathbb{E}_{\mathbf{x}}\mathbb{E}\left[2\left(\sum_{k<l} A_{kl}z_k z_l\right)\left(\sum_{k,l} A_{kl}\epsilon_k z_l\right)|\mathbf{x}\right] = 0. \tag{28}$$

**Derivation of** $\mathrm{Var}(\tilde{\mathbf{y}}_{p_1}^T\tilde{\mathbf{K}}_{\mathrm{x}}\tilde{\mathbf{y}}_{p_2})$**.**

Combining the above results, we obtain

$$\begin{aligned}\mathrm{Var}(\tilde{\mathbf{y}}_{p_1}^T\tilde{\mathbf{K}}_{\mathrm{x}}\tilde{\mathbf{y}}_{p_2}) =& \sum_{k<l} A_{kl}^2 s_k s_l(\mu_k\mu_l + \Sigma_{kl}) \\ &+ \frac{1}{2}\mathrm{tr}[(\mathbf{AS\Sigma S})^2] \\ &+ \sum_{k<l,k<u} A_{kl}A_{ku}s_k\mu_k s_l s_u \Sigma_{lu} + \sum_{k>l,k>u} A_{kl}A_{ku}s_k\mu_k s_l s_u \Sigma_{lu}.\end{aligned} \tag{29}$$

For convenience, define

$$\mathbf{\Omega} = \begin{bmatrix} \mathbf{\Omega}_{p_1} & \mathbf{\Omega}_{\mathrm{x}} \\ \mathbf{\Omega}_{\mathrm{x}}^T & \mathbf{\Omega}_{p_2} \end{bmatrix} = \mathbf{S\Sigma S}, \tag{30}$$

where

$$\mathbf{\Omega}_{p_1} = \tau_{p_1}^2\tilde{\mathbf{K}}_{p_1} + \sigma_{p_1}^2\tilde{\mathbf{I}}_n, \tag{31}$$

$$\mathbf{\Omega}_{p_2} = \tau_{p_2}^2\tilde{\mathbf{K}}_{p_2} + \sigma_{p_2}^2\tilde{\mathbf{I}}_n, \tag{32}$$

$$\mathbf{\Omega}_{\mathrm{x}} = \delta\tilde{\mathbf{K}}_{\mathrm{x}}. \tag{33}$$

Recall that $\mathbf{A} = \begin{bmatrix} \mathbf{0} & \tilde{\mathbf{K}}_{\mathrm{x}} \\ \tilde{\mathbf{K}}_{\mathrm{x}}^T & \mathbf{0} \end{bmatrix}$. Let $i, j, t \in \{1, \cdots, n\}$. Then the terms in $\mathrm{Var}(\tilde{\mathbf{y}}_{p_1}^T\tilde{\mathbf{K}}_{\mathrm{x}}\tilde{\mathbf{y}}_{p_2})$ can be expressed as follows.

$$\begin{aligned}\sum_{k<l} A_{kl}^2 s_k s_l(\mu_k\mu_l + \Sigma_{kl}) =& \sum_{i,j}\tilde{\mathbf{K}}_{\mathrm{x},ij}^2 s_i s_j(\mu_{p_1}\mu_{p_2} + \delta\mathbf{K}_{\mathrm{x},ij}) \\ =& \mu_{p_1}\mu_{p_2}\sum_{i,j}\tilde{\mathbf{K}}_{\mathrm{x},ij}^2 s_i s_j + \delta\sum_{i,j}\tilde{\mathbf{K}}_{\mathrm{x},ij}^3,\end{aligned} \tag{34}$$

$$\begin{aligned}\frac{1}{2}\mathrm{tr}[(\mathbf{AS\Sigma S})^2] =& \frac{1}{2}\mathrm{tr}[(\mathbf{A\Omega})^2] \\ =& \delta^2\mathrm{tr}[(\tilde{\mathbf{K}}_{\mathrm{x}}^T\tilde{\mathbf{K}}_{\mathrm{x}})^2] + \tau_{p_1}^2\tau_{p_2}^2\mathrm{tr}[\tilde{\mathbf{K}}_{\mathrm{x}}^T\tilde{\mathbf{K}}_{p_1}\tilde{\mathbf{K}}_{\mathrm{x}}\tilde{\mathbf{K}}_{p_2}] \\ &+ \tau_{p_1}^2\sigma_{p_2}^2\mathrm{tr}[\tilde{\mathbf{K}}_{\mathrm{x}}^T\tilde{\mathbf{K}}_{p_1}\tilde{\mathbf{K}}_{\mathrm{x}}\tilde{\mathbf{I}}] + \tau_{p_2}^2\sigma_{p_1}^2\mathrm{tr}[\tilde{\mathbf{K}}_{\mathrm{x}}\tilde{\mathbf{K}}_{p_2}\tilde{\mathbf{K}}_{\mathrm{x}}^T\tilde{\mathbf{I}}] \\ &+ \sigma_{p_1}^2\sigma_{p_2}^2\mathrm{tr}[\tilde{\mathbf{K}}_{\mathrm{x}}^T\tilde{\mathbf{I}}\tilde{\mathbf{K}}_{\mathrm{x}}\tilde{\mathbf{I}}],\end{aligned} \tag{35}$$

$$\begin{aligned}&\sum_{k<l,k<u} A_{kl}A_{ku}s_k\mu_k s_l s_u\Sigma_{lu} + \sum_{k>l,k>u} A_{kl}A_{ku}s_k\mu_k s_l s_u\Sigma_{lu} \\ =& \sum_{i,j,t}\tilde{\mathbf{K}}_{\mathrm{x},ij}\tilde{\mathbf{K}}_{\mathrm{x},it}s_i\mu_{p_1}s_j s_t(\tau_{p_2}^2\mathbf{K}_{p_2,jt} + \sigma_{p_2}^2\mathbf{I}_{jt}) + \sum_{i,j,t}\tilde{\mathbf{K}}_{\mathrm{x},ij}^T\tilde{\mathbf{K}}_{\mathrm{x},it}^T s_i\mu_{p_2}s_j s_t(\tau_{p_1}^2\mathbf{K}_{p_1,jt} + \sigma_{p_1}^2\mathbf{I}_{jt}) \\ =& \mu_{p_1}\tau_{p_2}^2\sum_{i,j,t}\tilde{\mathbf{K}}_{\mathrm{x},ij}\tilde{\mathbf{K}}_{\mathrm{x},it}s_i\tilde{\mathbf{K}}_{p_2,jt} + \mu_{p_1}\sigma_{p_2}^2\sum_{i,j,t}\tilde{\mathbf{K}}_{\mathrm{x},ij}\tilde{\mathbf{K}}_{\mathrm{x},it}s_i\tilde{\mathbf{I}}_{jt} \\ &+ \mu_{p_2}\tau_{p_1}^2\sum_{i,j,t}\tilde{\mathbf{K}}_{\mathrm{x},ij}^T\tilde{\mathbf{K}}_{\mathrm{x},it}^T s_i\tilde{\mathbf{K}}_{p_1,jt} + \mu_{p_2}\sigma_{p_1}^2\sum_{i,j,t}\tilde{\mathbf{K}}_{\mathrm{x},ij}^T\tilde{\mathbf{K}}_{\mathrm{x},it}^T s_i\tilde{\mathbf{I}}_{jt}.\end{aligned} \tag{36}$$

Collecting all terms, we have

$$\begin{aligned}\mathrm{Var}(\tilde{\mathbf{y}}_{p_1}^T\tilde{\mathbf{K}}_{\mathrm{x}}\tilde{\mathbf{y}}_{p_2}) =& \mu_{p_1}\mu_{p_2}\sum_{i,j}\tilde{\mathbf{K}}_{\mathrm{x},ij}^2 s_i s_j + \delta\sum_{i,j}\tilde{\mathbf{K}}_{\mathrm{x},ij}^3 \\ &+ \delta^2\mathrm{tr}[(\tilde{\mathbf{K}}_{\mathrm{x}}^T\tilde{\mathbf{K}}_{\mathrm{x}})^2] + \tau_{p_1}^2\tau_{p_2}^2\mathrm{tr}[\tilde{\mathbf{K}}_{\mathrm{x}}^T\tilde{\mathbf{K}}_{p_1}\tilde{\mathbf{K}}_{\mathrm{x}}\tilde{\mathbf{K}}_{p_2}] \\ &+ \tau_{p_1}^2\sigma_{p_2}^2\mathrm{tr}[\tilde{\mathbf{K}}_{\mathrm{x}}^T\tilde{\mathbf{K}}_{p_1}\tilde{\mathbf{K}}_{\mathrm{x}}\tilde{\mathbf{I}}] + \tau_{p_2}^2\sigma_{p_1}^2\mathrm{tr}[\tilde{\mathbf{K}}_{\mathrm{x}}\tilde{\mathbf{K}}_{p_2}\tilde{\mathbf{K}}_{\mathrm{x}}^T\tilde{\mathbf{I}}] + \sigma_{p_1}^2\sigma_{p_2}^2\mathrm{tr}[\tilde{\mathbf{K}}_{\mathrm{x}}^T\tilde{\mathbf{I}}\tilde{\mathbf{K}}_{\mathrm{x}}\tilde{\mathbf{I}}] \\ &+ \mu_{p_1}\tau_{p_2}^2\sum_{i,j,t}\tilde{\mathbf{K}}_{\mathrm{x},ij}\tilde{\mathbf{K}}_{\mathrm{x},it}s_i\tilde{\mathbf{K}}_{p_2,jt} + \mu_{p_1}\sigma_{p_2}^2\sum_{i,j,t}\tilde{\mathbf{K}}_{\mathrm{x},ij}\tilde{\mathbf{K}}_{\mathrm{x},it}s_i\tilde{\mathbf{I}}_{jt} \\ &+ \mu_{p_2}\tau_{p_1}^2\sum_{i,j,t}\tilde{\mathbf{K}}_{\mathrm{x},ij}^T\tilde{\mathbf{K}}_{\mathrm{x},it}^T s_i\tilde{\mathbf{K}}_{p_1,jt} + \mu_{p_2}\sigma_{p_1}^2\sum_{i,j,t}\tilde{\mathbf{K}}_{\mathrm{x},ij}^T\tilde{\mathbf{K}}_{\mathrm{x},it}^T s_i\tilde{\mathbf{I}}_{jt}. \end{aligned} \tag{37}$$

Finally,

$$\mathrm{Var}(\hat{\delta}) = \frac{\mathrm{Var}(\tilde{\mathbf{y}}_{p_1}^T\tilde{\mathbf{K}}_{\mathrm{x}}\tilde{\mathbf{y}}_{p_2})}{[\mathrm{tr}(\tilde{\mathbf{K}}_{\mathrm{x}}^T\tilde{\mathbf{K}}_{\mathrm{x}})]^2}. \tag{38}$$

### Supplementary Note 3: Selection of the spatial kernel

Following the SPARK method for univariate expression modeling in spatial transcriptomics [1], we consider multiple candidate spatial kernels. In the bivariate analyses, for each gene of interest, we select the optimal spatial kernel that provides the most accurate marginal fit to the expression data. Specifically, for gene $q$, we assess the quality of its marginal fit under the $k$-th spatial kernel $\mathbf{K}_k$ using the fitting accuracy score $F$:

$$\begin{aligned} F &= \|\tilde{\mathbf{y}}_q\tilde{\mathbf{y}}_q^T - \tilde{\mathbf{M}}_q\|_F, \\ \tilde{\mathbf{M}}_q &= \mathrm{diag}(\tilde{\boldsymbol{\mu}}_q) + \tau_q^2\tilde{\mathbf{K}}_k + \sigma_q^2\tilde{\mathbf{I}}. \end{aligned} \tag{39}$$

To compute $F$, we expand its expression to:

$$\begin{aligned} F =& \mathrm{tr}\{[\tilde{\mathbf{y}}_q\tilde{\mathbf{y}}_q^T - \mathrm{diag}(\tilde{\boldsymbol{\mu}}_q) - \tau_q^2\tilde{\mathbf{K}}_k - \sigma_q^2\tilde{\mathbf{I}}][\tilde{\mathbf{y}}_q\tilde{\mathbf{y}}_q^T - \mathrm{diag}(\tilde{\boldsymbol{\mu}}_q) - \tau_q^2\tilde{\mathbf{K}}_k - \sigma_q^2\tilde{\mathbf{I}}]^T\} \\ =& (\tau_q^2)^2\mathrm{tr}(\tilde{\mathbf{K}}_k^2) + (\sigma_q^2)^2\mathrm{tr}(\tilde{\mathbf{I}}^2) + 2\tau_q^2\sigma_q^2\mathrm{tr}(\tilde{\mathbf{K}}_k\tilde{\mathbf{I}}) \\ &- 2\tau_q^2\{\tilde{\mathbf{y}}_q^T\tilde{\mathbf{K}}_k\tilde{\mathbf{y}}_q - \mathrm{tr}[\mathrm{diag}(\tilde{\boldsymbol{\mu}}_q)\tilde{\mathbf{K}}_k]\} - 2\sigma_q^2\{\tilde{\mathbf{y}}_q^T\tilde{\mathbf{I}}\tilde{\mathbf{y}}_q - \mathrm{tr}[\mathrm{diag}(\tilde{\boldsymbol{\mu}}_q)\tilde{\mathbf{I}}]\} \\ &+ \mathrm{tr}\{[\tilde{\mathbf{y}}_q\tilde{\mathbf{y}}_q^T - \mathrm{diag}(\tilde{\boldsymbol{\mu}}_q)][\tilde{\mathbf{y}}_q\tilde{\mathbf{y}}_q^T - \mathrm{diag}(\tilde{\boldsymbol{\mu}}_q)]^T\}. \end{aligned} \tag{40}$$

The last term in Supplementary Eq. (40) is constant across spatial kernels, and the remaining terms are already included in Eq. (12). Therefore, no additional matrix computations are required, making the selection of the optimal spatial kernel computationally efficient.

### Supplementary Figures

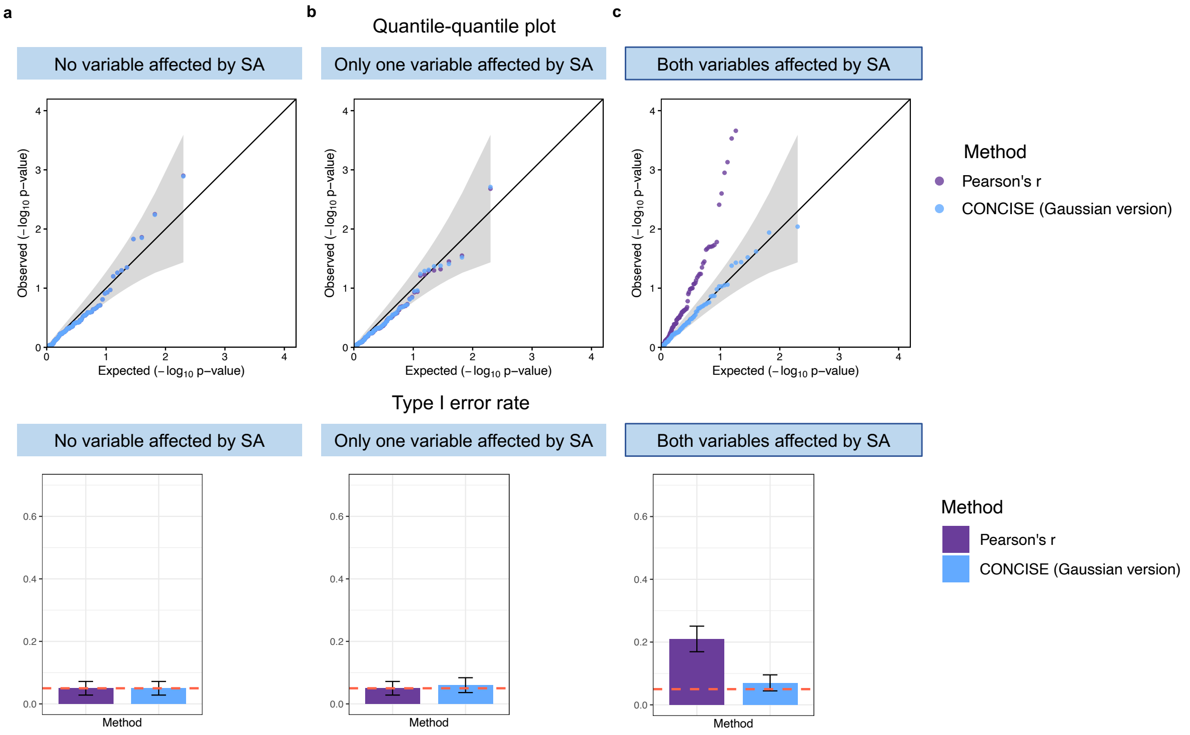


Supplementary Fig. 1: ***p*-value calibration and type I error control in spatial gene-gene co-expression analysis.** Two independent variables were simulated under three null scenarios: neither variable (**a**), only one variable (**b**), or both variables (**c**) exhibiting spatial autocorrelation (SA). Spatial co-expression was assessed using the Gaussian variant of CONCISE and Pearson's correlation. For each scenario, the top panel shows quantile-quantile plots of the resulting *p*-values, and the bottom panel summarizes the corresponding type I error rates. Results are based on 100 simulation experiments.

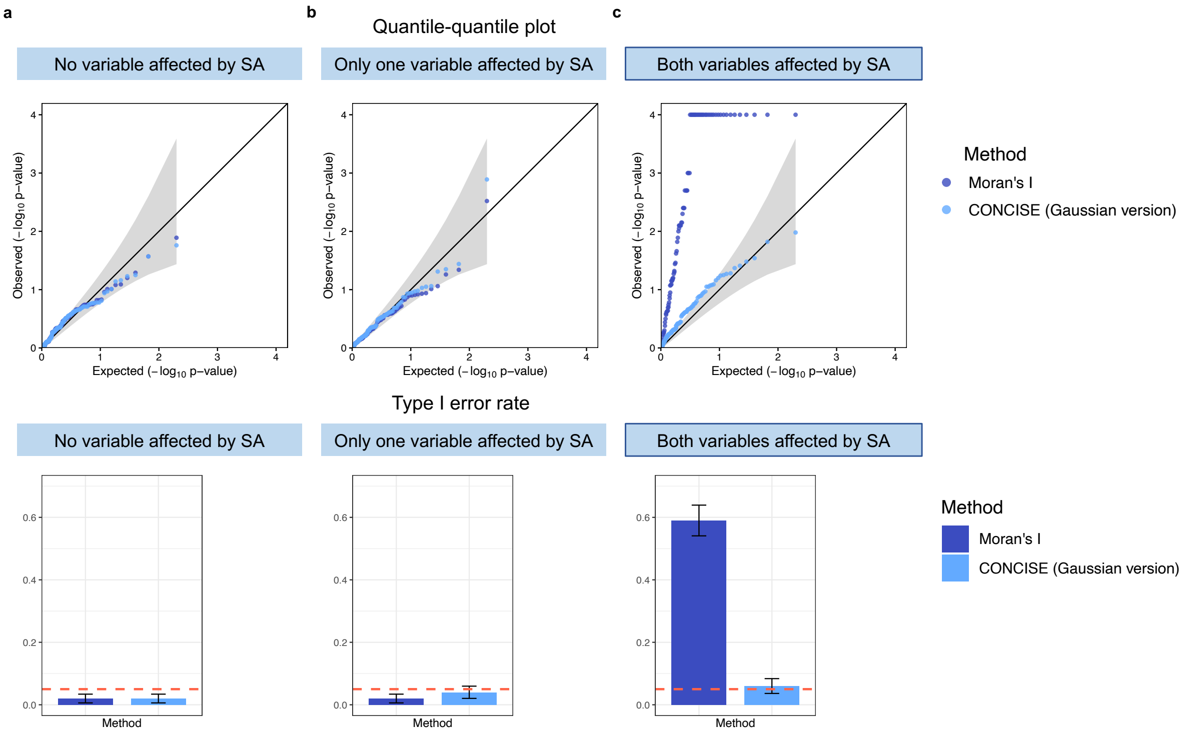


Supplementary Fig. 2: **$p$-value calibration and type I error control in spatially constrained co-expression analysis.** Two independent variables were simulated under three null scenarios: neither variable (**a**), only one variable (**b**), or both variables (**c**) exhibiting spatial autocorrelation (SA). Spatially constrained co-expression was assessed using bivariate Moran's I with permutation testing and the Gaussian variant of CONCISE. For each scenario, the top panel shows quantile-quantile plots of the resulting $p$-values, and the bottom panel summarizes the corresponding type I error rates. Results are based on 100 simulation experiments.

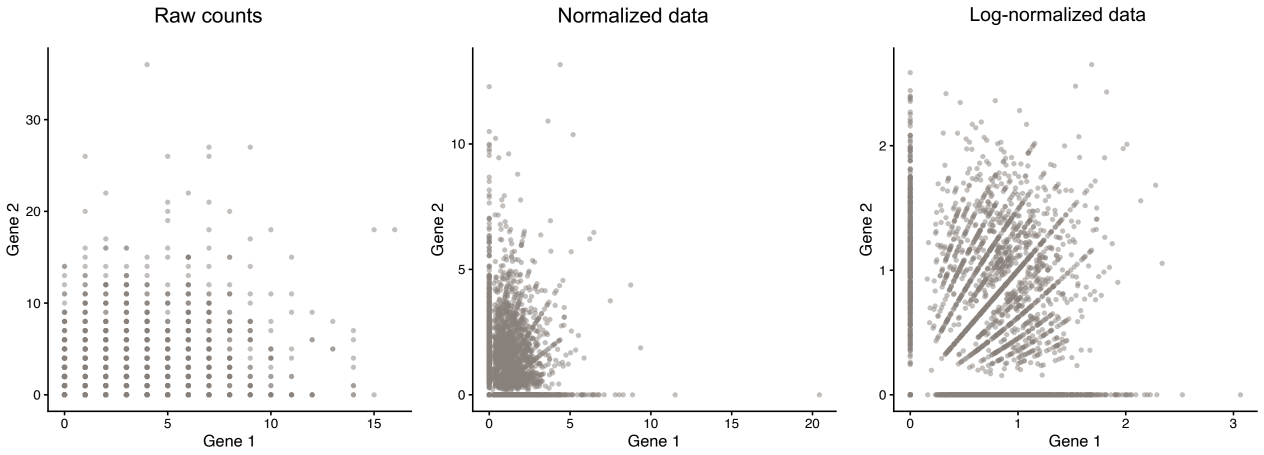


Supplementary Fig. 3: **Expression patterns of an independent gene pair in the real data-based permutation study, shown using raw counts, normalized counts, and log-normalized counts.**

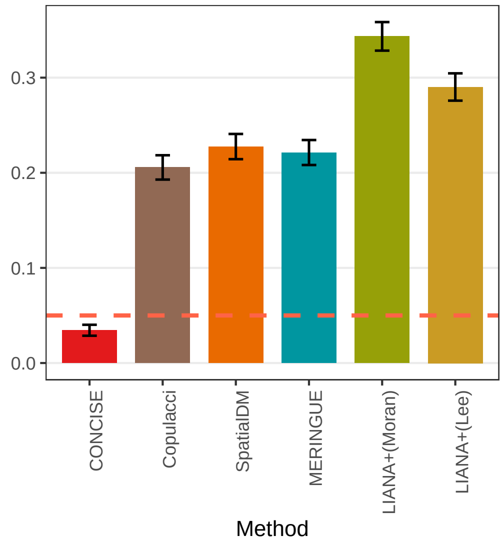


Supplementary Fig. 4: **Comparison of type I error rates across methods in simulations of L-R pairs parameterized using the breast cancer dataset.** Type I error rates were evaluated at a significance threshold of $p$-value=0.05.

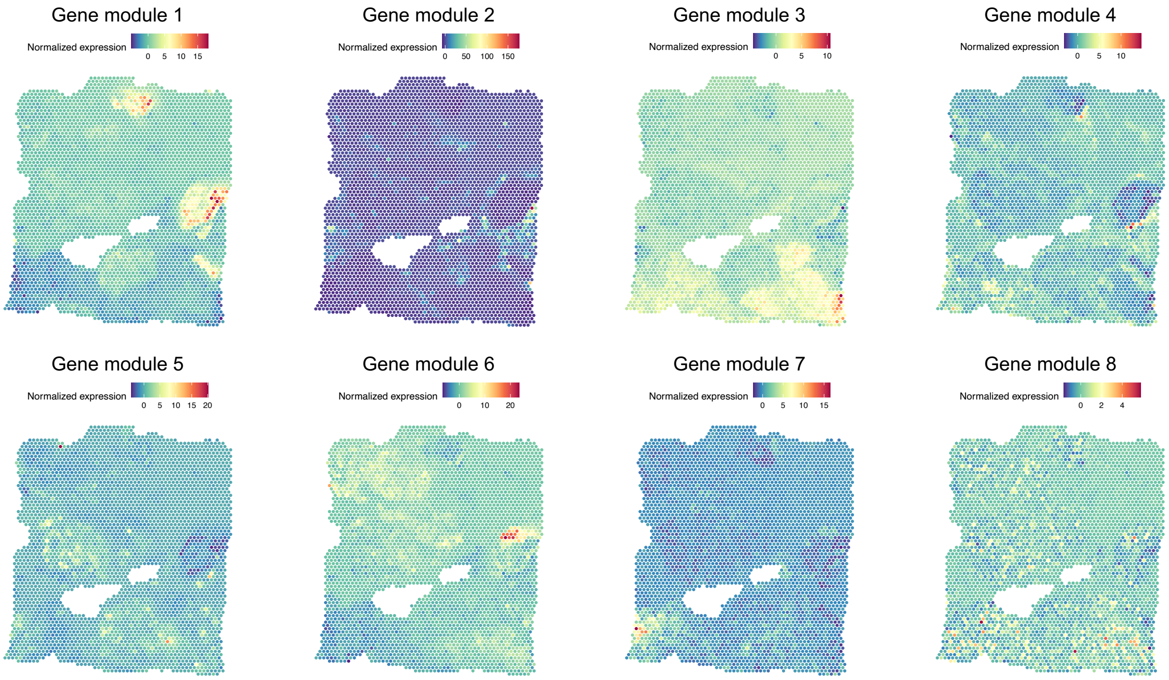


Supplementary Fig. 5: **Spatial patterns of gene modules identified from the CONCISE's inferred gene-gene co-expression network in the breast cancer dataset.**

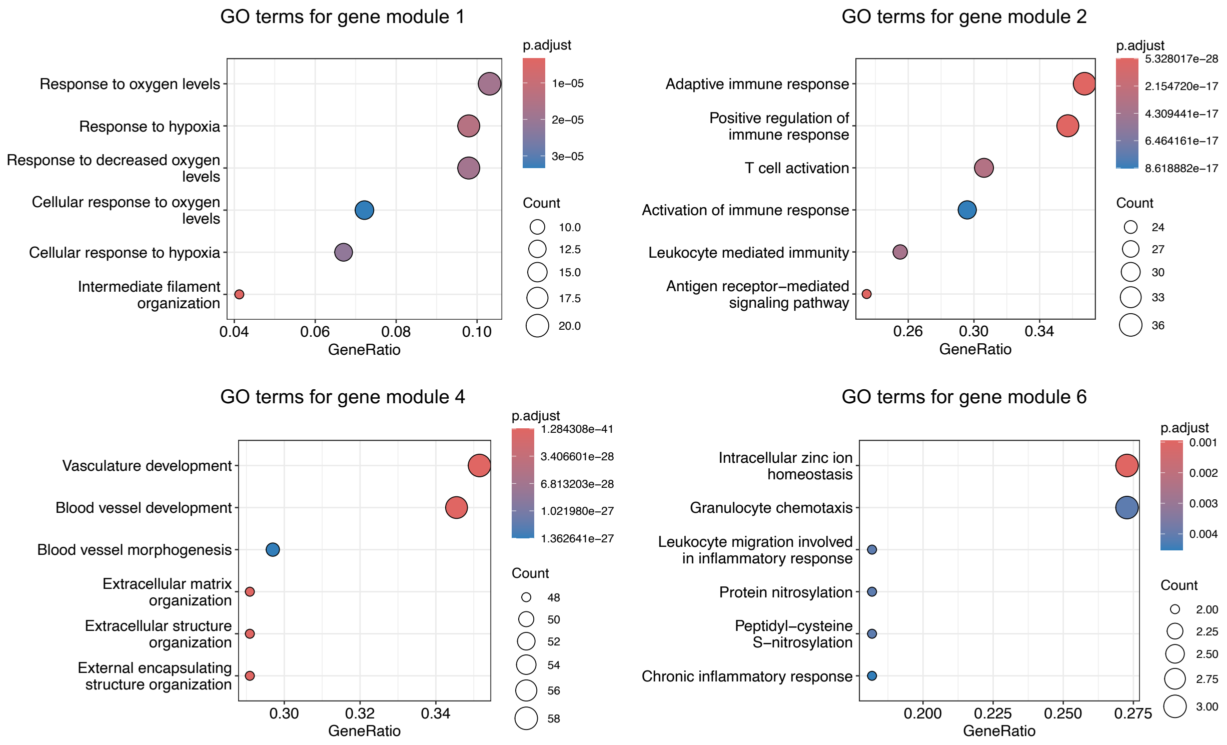


Supplementary Fig. 6: **Significantly enriched GO terms for gene modules spatially localized near the tumor region.**

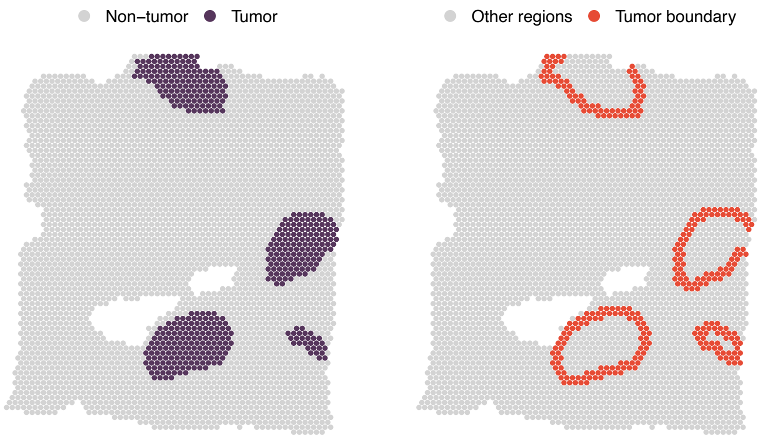


Supplementary Fig. 7: **Tumor core and tumor boundary regions in the Visium breast cancer dataset.**

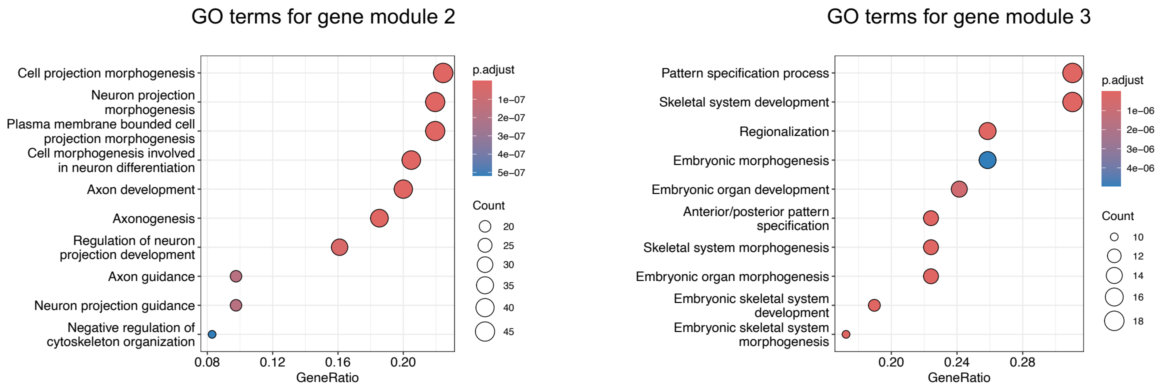


Supplementary Fig. 8: **GO enrichment analysis of gene modules 2 and 3 identified from the CONCISE's inferred spatial gene-gene co-expression network in the Stereo-seq mouse embryo dataset.** Gene module 2 is enriched for neuronal development-related processes, whereas gene module 3 is enriched for embryonic skeletal system development and embryonic patterning.

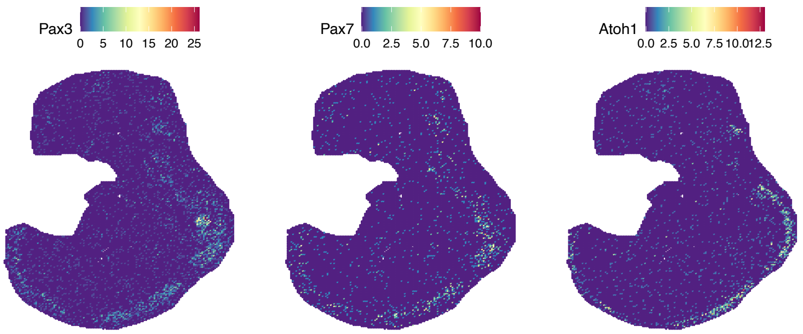


Supplementary Fig. 9: **Spatial expression patterns of the dorsal neural tube-related marker genes *Pax3*, *Pax7*, and *Atoh1*.**

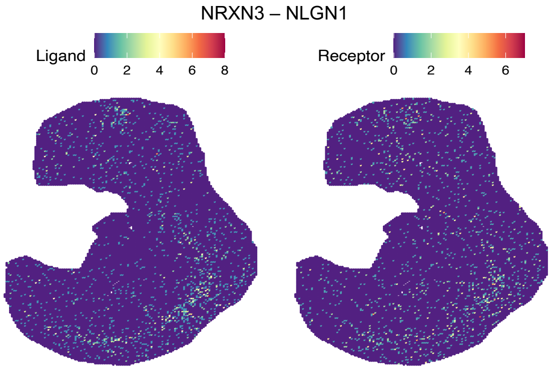


Supplementary Fig. 10: **Spatial expression patterns of the NRXN3-NLGN1 interaction.**

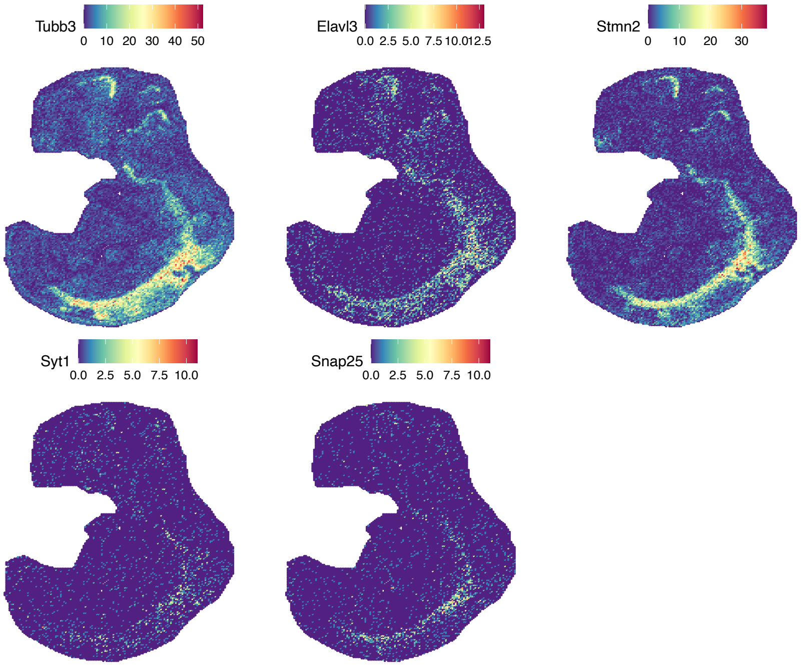


Supplementary Fig. 11: **Spatial expression patterns of neuronal marker genes and genes involved in synaptic function.** Neuronal marker genes include *Tubb3*, *Elavl3*, and *Stmn2*. Genes involved in synaptic function include *Syt1* and *Snap25*.

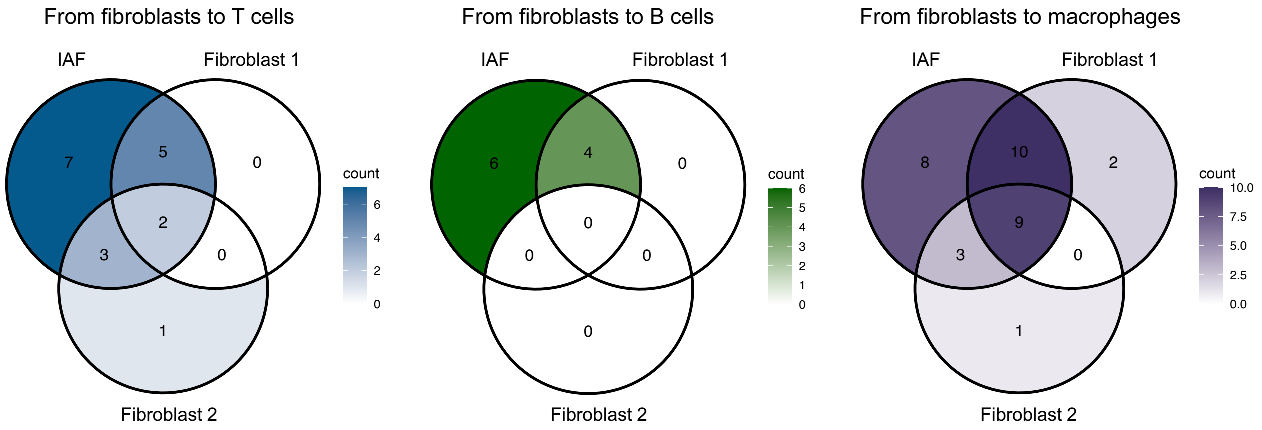


Supplementary Fig. 12: **Overlap of significant LRIs from fibroblast populations to immune cells.** Venn diagrams showing significant LRIs identified between fibroblast populations and T cells (left), B cells (middle), and macrophages (right). Inflammation-associated fibroblasts (IAFs) exhibit a substantially larger number of unique interactions with all three immune cell populations than homeostatic fibroblasts, indicating enhanced communication activity associated with the inflammatory fibroblast state.

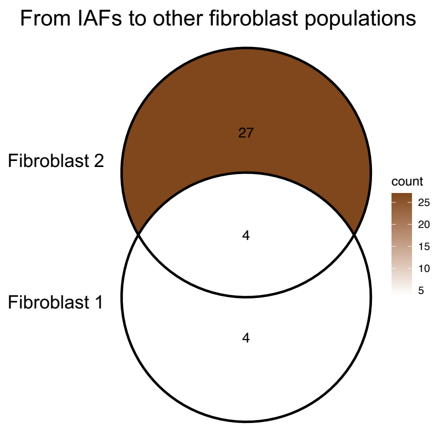


Supplementary Fig. 13: **Overlap of significant LRIs from inflammation-associated fibroblasts to homeostatic fibroblast populations.** Venn diagram showing the overlap of significant LRIs inferred from IAFs to fibroblast 1 and fibroblast 2. Shared and population-specific interactions are indicated.

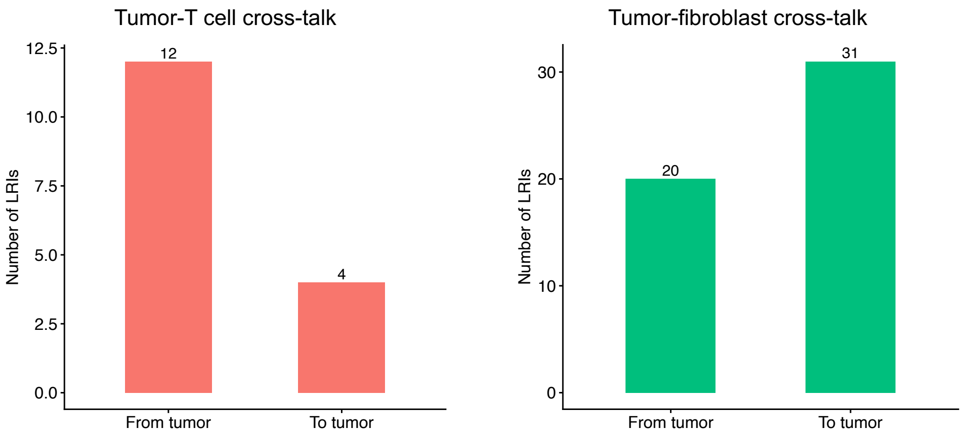


Supplementary Fig. 14: **Numbers of significant LRIs between tumor cells and T cells or fibroblasts.** Significant LRIs are grouped according to directionality, including interactions originating from tumor cells and interactions directed toward tumor cells.
